# SOX9-positive pituitary stem cells differ according to their position in the gland and maintenance of their progeny depends on context

**DOI:** 10.1101/2022.11.07.515440

**Authors:** Karine Rizzoti, Probir Chakravarty, Daniel Sheridan, Robin Lovell-Badge

**Affiliations:** Laboratory of Stem Cell Biology and Developmental Genetics, The Francis Crick Institute; Bioinformatics core, The Francis Crick Institute, London NW1 1AT, UK

## Abstract

Stem cell (SC) differentiation and maintenance of resultant progeny underlie cell-turnover in many organs, but it is difficult to pinpoint the contribution of either process. In the pituitary, a central regulator of endocrine axes, adult SCs undergo activation following target organ ablation, providing a well-characterized paradigm to study an adaptative response in a multi-organ system. Here we used single cell technologies to characterize SC heterogeneity and mobilization together with lineage tracing. We show that SC differentiation occurs more frequently than thought previously. In adaptative conditions, differentiation increases and is more diverse than demonstrated by the lineage tracing experiments. Detailed examination of SC progeny suggests that maintenance of selected nascent cells underlies SC output, highlighting a trophic role for the microenvironment. Analyses of cell trajectories further predict pathways and potential new regulators. Our model provides a valuable system to study the influence of evolving states on the mechanisms of SC mobilization.

Teaser: Pituitary stem cells are diverse and differentiate more than thought but only selected progeny persist according to need.

## INTRODUCTION

The pituitary gland controls the three major endocrine axes, reproduction, stress and metabolism, and is involved in homeostasis; all under control of the hypothalamus. Its function requires modulation of hormonal output over short and long terms, and this is regulated at several levels, including rates and patterns of hormone secretion, synthesis and storage, and changes in endocrine cell numbers. Different mechanisms are known to underlie endocrine cell emergence: division of endocrine cells, differentiation from resident SCs and transdifferentiation, although the relevance of the latter is unclear. Under normal and stable conditions, it is thought that division from endocrine cell types is sufficient for the typical low cell turnover seen in the pituitary (*1*). This has been directly shown in corticotrophs using a genetic trick where cell division induces apoptosis of POMC expressing cells. A gradual loss of corticotrophs is observed in these pituitaries in support of the role of these endocrine cells in their own replacement (*2*). In agreement, long term lineage tracing of adult SOX2 and SOX9 +ve pituitary SCs in steady-state conditions suggests that SCs differentiate rarely and thus do not significantly participate in cell turnover (*3*) (*4*). Furthermore, adult ablation of the SCs does not appear to affect pituitary function (*5*). In contrast, when the regenerative or adaptative potential of the gland is stimulated, either by endocrine cell ablation (*6*) or target organ ablation (*3*) respectively, SC-derived new endocrine cells of the appropriate type emerge. In this latter paradigm, differentiation from SCs appears to be the main process underlying emergence of new corticotrophs after adrenalectomy (Ax) (*2*), providing a well characterized multi-organ system to explore mechanisms of SC activation.

Here, we explore the characteristics of pituitary SC fate acquisition using single cell approaches combined with lineage tracing analyses. We observe that SOX9 +ve pituitary SCs comprise several functionally distinct sub-populations, located in the ventral and dorsal part of the epithelium lining the pituitary cleft and in the anterior pituitary parenchyma. We then explored mechanisms underlying SC mobilization following target organ ablation through characterizing cell trajectories and gene regulatory networks (GRNs) potentially involved in cell fate acquisition. We observed that while fate acquisition of the targeted cell type (corticotrophs after Ax and gonadotrophs after gonadectomy) was efficiently induced, differentiation toward all endocrine cell types unexpectedly also took place. Examination of longer-term outcomes using lineage tracing further showed that only a proportion of the differentiated progeny is retained, specifically that of the targeted cell type. More precisely, we only observe SC-derived corticotrophs after Ax, and lactotrophs in females, following Selective Estrogen Receptor Modulator (SERM) treatment. Therefore, while pituitary SC differentiation appears both more frequent and diverse than previously thought, our results suggest that the selective maintenance of nascent cells, according to perceived pathophysiological need and likely sex-dependent hormonal cues, is an important regulatory step controlling their adaptative response. Our model thus allows characterisation of this response following pathophysiological manipulations that may well relate to normal life-changing events, such as puberty, pregnancy and lactation.

## RESULTS

### Distinct regulatory pathways govern AL and IL SC identity

The anterior lobes of three *Sox9^iresGFP/+^* adult male pituitaries were dissociated and the transcriptome of single GFP +ve cells analysed after fluorescence activated cell sorting (FACS), as previously described (*3*). After quality checks, 1993 cells were retained for further analysis; clustering was performed by generating UMAP plots. Three major clusters were identified (Fig.1A). Co-expression of *Sox2* and *Sox9* in clusters 0 and 2 (Fig.1B, Sup.Fig.1), representing 69% of the cells, suggested that these were likely to represent SCs. The presence of two SC clusters had previously been reported in whole pituitary datasets (*7*) (*8*). The remaining cluster (1), representing 24% of the dataset, comprised cells where *Sox*9 was detected, but where *Sox2* was greatly reduced or absent. Finally, cells each expressing a different hormone and the corresponding lineage specifying factors, were grouped within a single smaller cluster (3) representing 3% of the cells (Fig.1B, Sup.Fig.1). SOX9 is not expressed in any endocrine cell in quiescent conditions (*9*). However, we observed some rare SOX9;ACTH +ve cells, and more numerous GFP;ACTH +ve cells in *Sox9^iresGFP/+^*animals exclusively after activation of the SCs by pituitary target organ ablation (*3*) (Sup. Fig.2). This suggests that differentiating, and recently differentiated cells are present in the SOX9iresGFP fraction due to persistence of the GFP, which correlates with what has been suggested to happen in both human and mouse pituitaries, where hormone positive cells were included in the SC compartment of whole pituitary single cell RNAseq datasets (*10*). In addition, a small proportion of contaminating pituitary cells (estimated to represent 25% of cluster 3) is likely to be present (Sup.Fig.3). Therefore, most cells present in cluster 3 are likely to represent cells committed toward an endocrine fate, with the largest population (41% of cluster 3 cells) expressing *Prolactin*.

**Figure 1:**
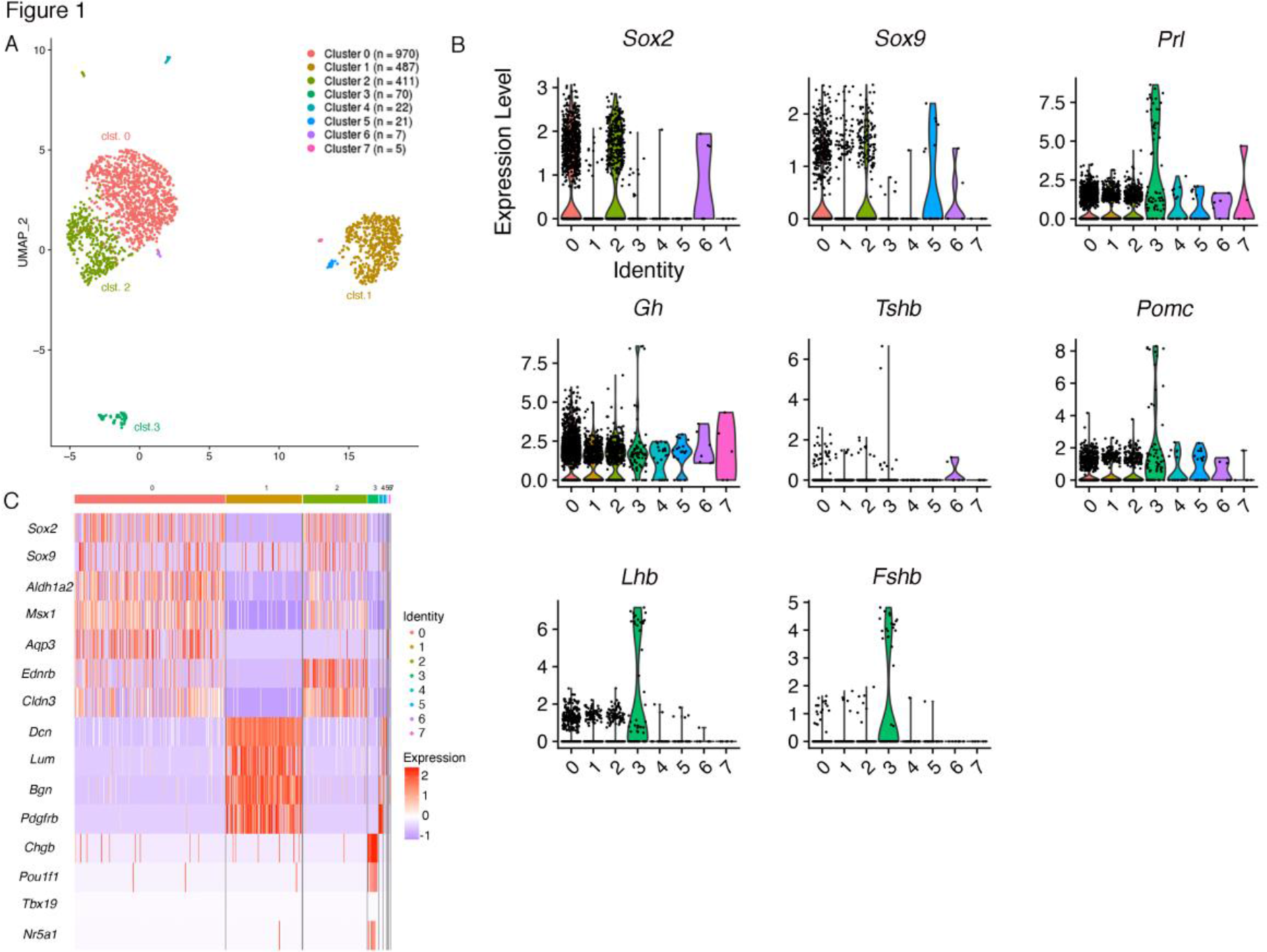
Single cell analysis of the *Sox9^iresGFP/+^* population in male pituitaries. A) UMAP clustering of *Sox9^iresGFP/+^* cells. B) Expression levels of *Sox2* and *Sox9* were examined to identify SCs (*9*) while genes encoding for hormones (*Gh*, *Prl*, *Tshβ*, *Lhβ*, *Fshβ*, *Pomc*) were examined to distinguish committed cells (*Pou1f1* for somatotrophs, lactotrophs and thyrotrophs. C) Heatmap displaying selected markers for the main clusters.

The two SC clusters were further explored. Analysis of differentially expressed genes (DEG) (Fig.1C, Sup.Fig.1B) shows that clusters 0 and 2 are relatively similar and are mostly distinguished by differences in levels of expression of specific genes. One notable exception is *Aqp3*, encoding for a water channel, which is specific for cluster 0. In contrast, *Aldh1a2*, encoding for an enzyme involved in retinoic acid (RA) synthesis, and the transcription factor *Msx1* were preferentially present in cluster 0, while *Ednrb*, encoding for the endothelin B receptor, and the cell adhesion molecule *Claudin3* were both expressed at higher levels in cluster 2. RA is required during pituitary morphogenesis (*11*) (*12*) and *Msx1* interacts with SOX2 for *Prop1* regulation (*13*), while nothing is known about *Ednrb* or *Claudin3* in pituitary SCs. Conservation of most of these markers in SCs from whole pituitary datasets (*7*) (*8*) suggests that our two SC clusters correlate with those observed by others. In the gland, expression of the five markers is restricted to SCs (Fig.2, Sup. Fig.4). In the epithelium lining the cleft, cluster 0 markers ALDH1a2 and MSX1 are expressed exclusively on the AL side (Fig.2 A,, Sup.Fig4B), while cluster 2 markers EDNRB and CLAUDIN-3 (Fig.2 C,D) are present on both sides. In the parenchyma ALDH1a2, MSX1 and CLAUDIN-3 co-localise with SOX2/9 while expression of EDNRB is not detectable. In situ hybridisation for *Ednrb* confirms a clear enrichment in cleft lining cells (Sup.Fig.5). AQP3, whose transcript is present in about half of cluster 0 cells, is exclusively detected in the parenchymal SCs, revealing further heterogeneity between cleft-lining and parenchymal populations. In addition, it is only expressed in a subset of SOX2;SOX9 positive parenchymal SCs (Fig.2B), demonstrating the existence of parenchymal subpopulations. Examination of the five markers in scRNAseq datasets from whole pituitaries (21) confirms that the strongest, and sometimes exclusive, expression is seen in SCs (Sup. Fig.4). All together, these results suggest that cluster 0 is enriched in anterior lobe SCs, while cluster 2 has a more generic pituitary SC identity.

**Figure 2:**
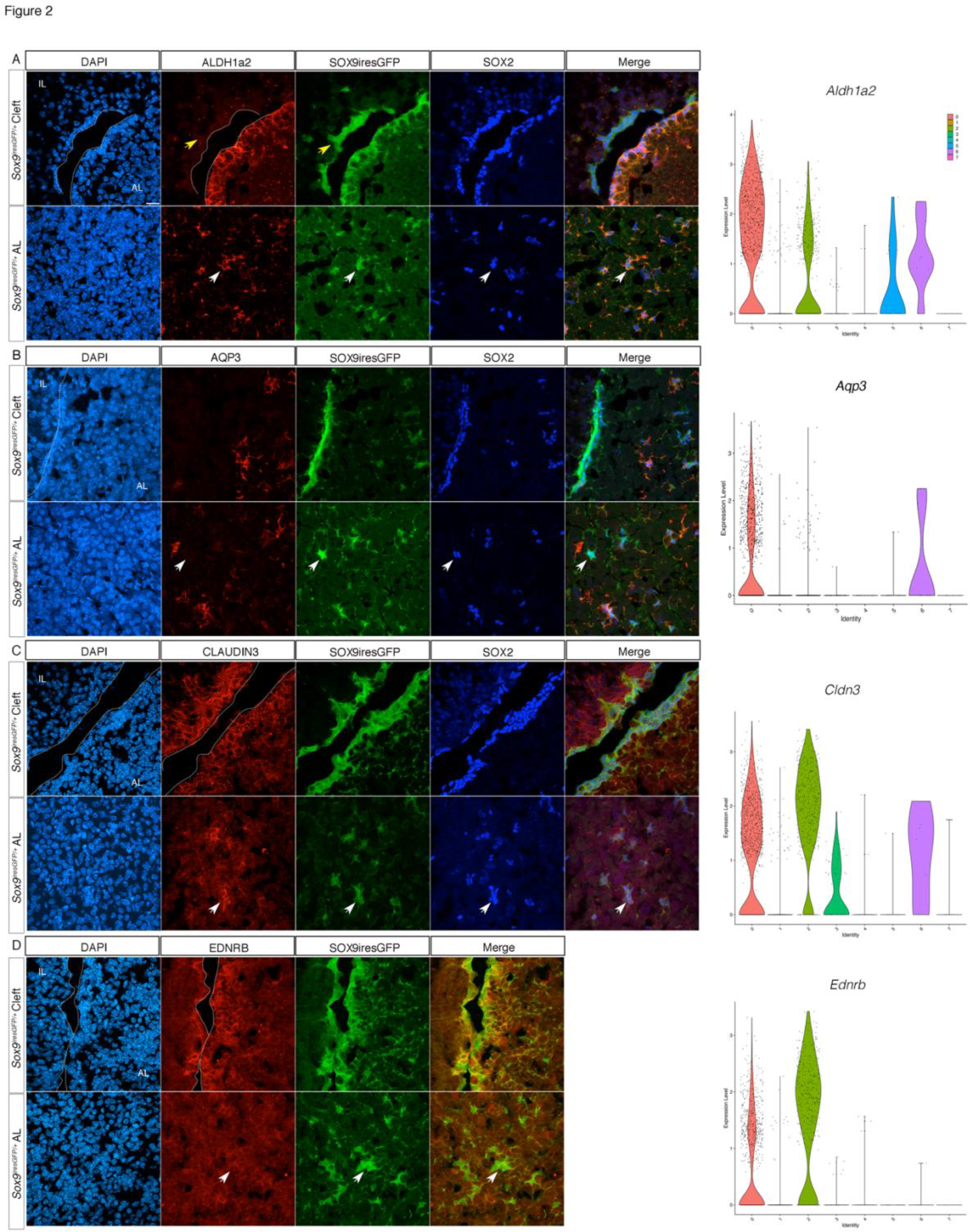
Differential expression of SC cluster markers. A) Co-immunofluorescence for ALDH1A2, GFP and SOX2 in a male SOX9iresGFP pituitary section. ALDH1A2 is exclusively expressed in SOX2;SOX9 AL SC (yellow arrow shows its absence from IL SC, and white arrow shows its expression in parenchymal SC). *Aldh1a2* violin plot shows enrichment in cluster 0. B) Co-immunofluorescence for AQP3, GFP and SOX2 in a male SOX9iresGFP pituitary section. AQP3 is expressed in 66% of SOX2 +ve parenchymal SC (white arrow shows SOX2;SOX9iresGFP parenchymal SC not expressing AQP3; counting was performed on n=3 animals). *Aqp3* violin plot shows enrichment in cluster 0. C) Co-immunofluorescence for CLAUDIN3, GFP and SOX2 in a male SOX9iresGFP pituitary section. CLAUDIN3 is exclusively expressed in SOX2;SOX9 SC. *Cldn3* UMAP shows enrichment in cluster 2. D) Co-immunofluorescence for EDNRB and GFP in a male SOX9iresGFP pituitary section. EDNRB is exclusively expressed in SOX9 cleft SC. *Ednrb* UMAP shows enrichment in cluster 2. Similar expression profiles were observed in females for all markers. The scale bar in A represents 10 μm for all panels. White arrows show absence of expression in parenchymal SC. AL, anterior lobe, IL intermediate lobe.

Pathway analysis in cluster 0 (Sup Table.1) highlighted activity of the NOTCH and WNT pathways and presence of Ephrin signalling, as previously reported in pituitary SCs (*14*) (*15*). Cluster 2 cells are characterised by expression of cytoskeleton remodelling and adhesion genes consistent with the EMT-like process required during pituitary progenitor cell fate acquisition (*16*). In conclusion, regionalisation of these two SC subpopulations is underlined by the expression of genes involved in the activity of specific regulatory pathways, such as retinoic acid signalling in AL SCs. Further investigation is required to understand what distinguishes AQP3 positive and negative parenchymal SCs.

Folliculo-stellate (FS) cells are specialised AL glial cells (*17*). These express S100b in the rat, but not in the mouse ((*18, 19*), Sup. Fig.6). To clarify their relationship with SCs, we examined expression of *Aldolase C*, a marker for rodent FS cells (*19*) and of *Slc152a*, which encodes a transporter responsible for the FS specific import of AMCA (*20*). Both are selectively present in SCs, in clusters 0 and 2 (Sup.Fig.6A). Furthermore, in a whole pituitary dataset, *Sox2* and *Aldoc* are exclusively present in the SC cluster (Sup.Fig.6B). All together these results confirm, as suggested earlier (*9, 21*) (*22, 23*), that in the mouse FS and SCs are overlapping populations.

In conclusion, single-cell analysis of the *Sox9^iresGFP/+^*pituitary fraction successfully resolved heterogeneity of this population, suggesting differential activity of signalling pathways, and clarified the relationship between SCs and FS cells.

### A novel population of parenchymal SOX9;PDGFRβ +ve cells

Cluster 1, representing 25% of the cells, is characterised by expression of PDGFRβ (Fig.1C, Fig.3E). *In situ*, PDGFRβ;SOX9 +ve cells are exclusively detected in the parenchyma. Expression of SOX9 in these double +ve cells is weaker than in SCs (Fig.1B, Fig.3A) and SOX2 is not detected (Fig.1B , Fig.3B). In contrast with the *in silico* analysis, only 8±1.62% SD of parenchymal SOX9 +ve cells express PDGFRβ in males (Fig.3C). This discrepancy could be due either to an over-representation of cluster 1 cells, because epithelial SCs are more difficult to recover than parenchymal cells, or to the difficulty in detecting the cells by immunofluorescence. We observe significantly fewer PDGFRβ;SOX9 +ve cells in females than in males (1.3±0.6% SD, p= 0.029, Fig.3C). PDGFRβ +ve pituitary cells are known to comprise a mural vascular network (*18*). While we did not attempt to fully characterize the SOX9;PDGFRβ population, we observe some SOX9;PDGFRβ cells in close contact with capillaries, displaying a pericyte like morphology (Fig.3D). In the embryo, vascular and mural cells have an extra-Rathke’s pouch origin, as they emerge from immigrating neural crest cells (*24, 25*). Because cluster 1 cells do not express the generic pituitary marker *Pitx1* (Fig.3E), we analysed a potential neural crest origin by performing WNT1-Cre lineage tracing. 79 ± 3.1% SEM of SOX9;PDGFRβ cells (n=130 cells from 4 males) derive from WNT1-Cre +ve cells (Fig3C, F) while in females only 47.5 ± 5.5% SEM of the cells derive from the neural crest (n= 57 cells from 3 females, p= 0.024). Therefore, we propose that cluster 1 comprises a sexually dimorphic mural network with a dual embryonic origin. The pituitary capillary network, which controls endocrine output, adapts according to the physiological context and this is particularly important for gonadotroph function (*26*). The sex differences we observe may be related to this aspect.

**Figure 3:**
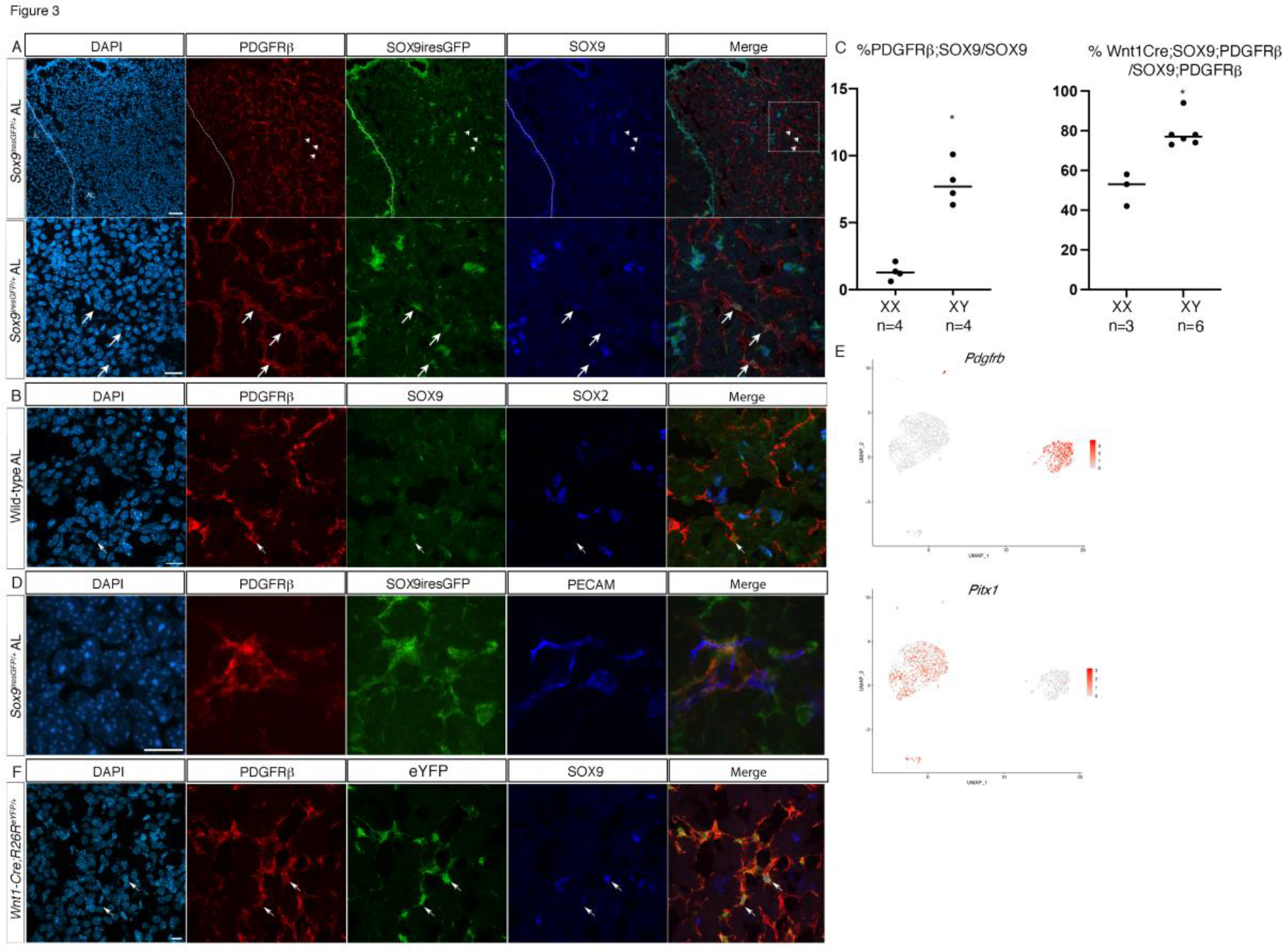
Characterisation of a SOX9;PDGFRβ +ve AL cell population. A) Co-immunofluorescence for PDGFRβ, GFP and SOX9 in a SOX9iresGFP male AL section. Triple +ve cells are indicated (arrows). The lower panels represent the magnification of the region indicated on the merge panel above. The dotted line indicates the pituitary cleft. B) Co-immunofluorescence for PDGFRβ, GFP and SOX2 in AL. A SOX9;PDGFRβ +ve cell which is negative for SOX2 (arrow). C) Percentage of GFP;SOX9;PDGFRβ/GFP;SOX9 +ve cells in *Sox9iresGFP* male and female AL sections (average 500 SOX9 cells counted/animal, p = 0.029). In SOX9iresGFP pituitary sections, parenchymal SOX9; GFP were captured (Sup.Table.4). Amongst these, the percentage of those also positive for PDGFRβ cells was determined. Percentage of SOX9;PDGFRβ +ve cells of neural crest origin (p = 0.024; Mann Whitney test). In Wnt1Cre;R26ReYFP pituitary sections, Sox9;PDGFRb cells were captured. Amongst these, the percentage of those also positive for Wnt1Cre;ReYFP cells was determined (Sup.Table.4). D) Co-immunofluorescence for PDGFRβ, GFP and PECAM in a Sox9iresGFP AL section showing a SOX9;PDGFRβ +ve cell resembling a pericyte bridging PECAM +ve capillaries. E) UMAP for *Pitx1* and *Pdgfrβ* showing that *Pdgfrβ* +ve cluster 1 cells do not express *Pitx1*. F) Co-immunofluorescence for PDGFRβ, eYFP and SOX9 in a *Wnt1-Cre;R26R^eYFP^* AL section. Triple +ve cells are shown (arrows). The scale bars represent 50 μm and 20 μm for A (upper and lower panels respectively) and 10 μm for D and F.

### All endocrine cell types are present in the *Sox9^iresGFP/+^* +ve compartment after SC mobilization

We then performed sc-RNAseq on activated SCs. Target organ ablation induces a transient proliferative wave in the gland, mostly affecting SCs, which can then give rise to new endocrine cells (*2, 3, 27*). We analysed the transcriptome of *Sox9^iresGFP/+^*male cells 4 days after removing the adrenals, the testes or both. We integrated these datasets with that obtained from unchallenged animals and obtained a similar clustering of SCs (Fig.1, Fig.4A,B, Sup. Fig.7A). A pairwise comparison for proportion test limited to SC clusters demonstrates a significant increase in proliferative cells after target organ ablation (Sup.Fig.7B), as previously shown (*3*). We also observe the emergence of distinct hormone +ve clusters representing in total 25 to 30% of all cells (Fig.4A,B) compared to 3.5% in the unchallenged dataset, suggesting increased differentiation. Target organ ablation is associated with increased secretion, and sometimes cell numbers, of the endocrine population regulating the resected organ. However, regardless of the glands removed, all endocrine cell lineages emerge in the Sox9iresGFP +ve fraction. While we expected *Pomc* +ve cells to prominently feature after Ax, these were outnumbered by *Prl* and *LHb*;*FSHb* +ve cells (Sup. Fig. 7C). Similarly, gonadectomies were expected to trigger commitment toward the gonadotroph cell fate, but *Gh* +ve cells were mostly observed. To characterise these *Sox9^iresGFP/+^*;hormone +ve cells we compared them with endocrine cells by integrating our datasets with one from whole pituitaries (*28*). *Sox9^iresGFP/+^*;hormone +ve cells cluster with endocrine cells (Fig.4C, D). Therefore, they represent differentiated cells, at least transcriptionally. In conclusion, target organ ablation induces differentiation of SCs, but there is an unexpected lack of specificity in the endocrine cell types produced, with all hormonal markers being up-regulated.

**Figure 4:**
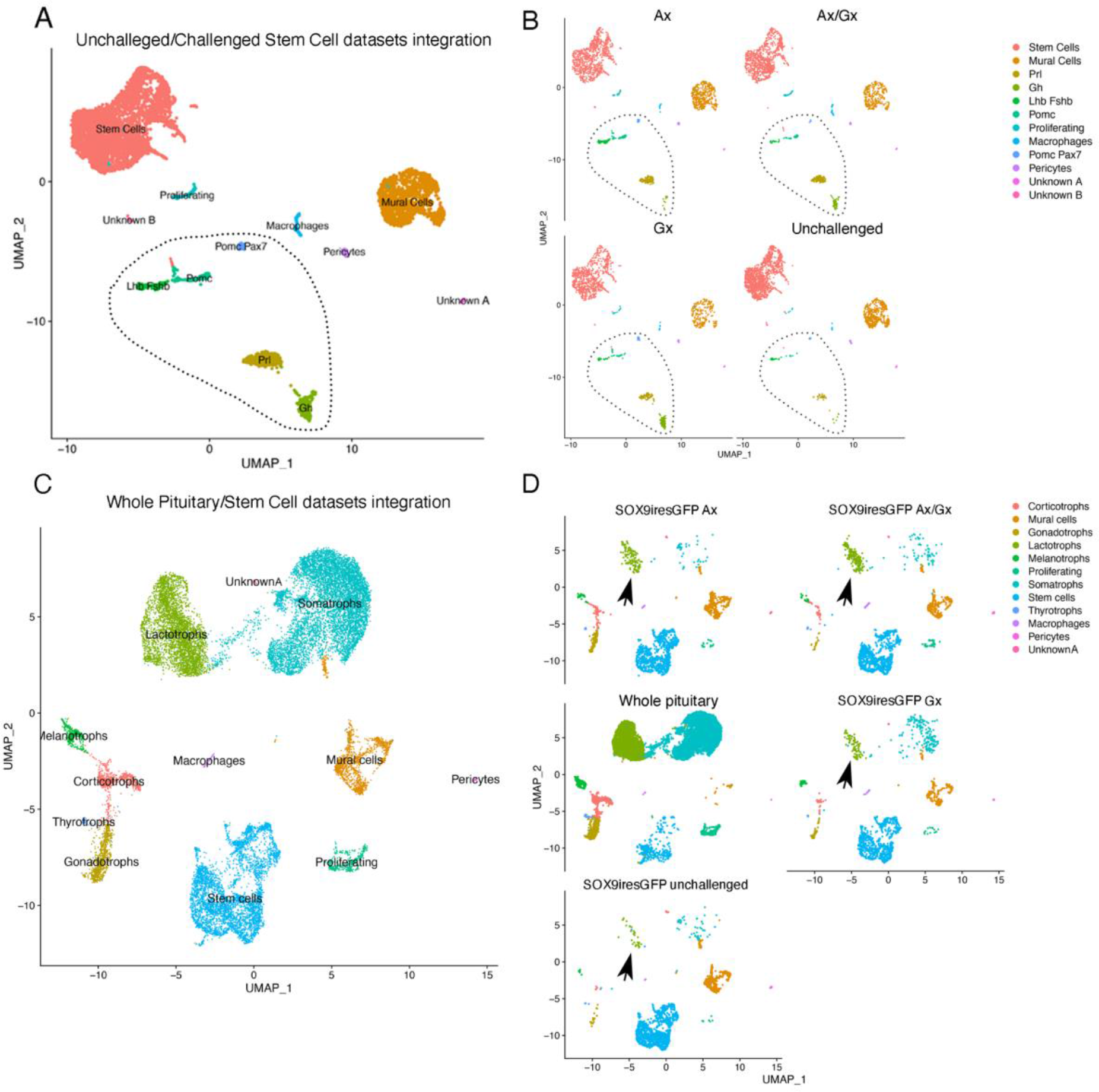
Target organ ablation induces differentiation toward multiple endocrine lineages. A) UMAP clustering for integrated datasets from sorted *Sox9^iresGFP/+^*cells from unchallenged, adrenalectomized (1915 cells), gonadectomized (1546 cells) and both adrenalectomized and gonadectomized (1880 cells) animals, 4 days after surgery. B) UMAP clustering split by experiment showing the contribution of cells originating from each dataset. Cells expressing all pituitary hormones are present (dotted lines) in all conditions. C) UMAP clustering for integrated datasets from sorted *Sox9^iresGFP/+^* cells and a whole pituitary dataset (*28*). D) UMAP clustering split by experiment show that hormone +ve cells present in the *Sox9^iresGFP/+^* cells cluster with endocrine cells (black arrow pointing to lactotrophs).

### Analyses of cell trajectories after Ax

We decided to focus on Ax because it represents the most efficient stimulus of SC differentiation after target organ ablation (*3*). We added a new time point at day 2 after Ax and a sham control. These new datasets were then integrated (Fig.5A) with our initial dataset (Fig.1) and the one performed 4 days after Ax (Fig.4). UMAP clustering indicated that SCs were present in four distinct clusters (Sup. Fig.4). In addition to the *Ednrb* and *Aldh1a2* clusters, two SC clusters (SC3 and SC4) emerged (Fig.5A). These were mostly but not exclusively populated by cells from the new datasets (Sup. Fig.8). While more cells were sequenced in these new datasets (up to 3 times more), we used a different chemistry (see material and methods) which, despite Seurat batch correction tools, may explain detection of more genes and cells resulting in emergence of these clusters. SC3 was characterised by higher expression of markers also present in the *Ednrb* and *Aldh1a2* clusters. SC4, mostly derived from the Ax+48h00 dataset, showed co-expression of SC and endocrine markers suggesting that it represents cells in a transitioning phase (Sup. Fig.9). Finally, endocrine cell clusters were detected (Sup. Fig.9), as observed previously (Fig.4).

**Figure 5:**
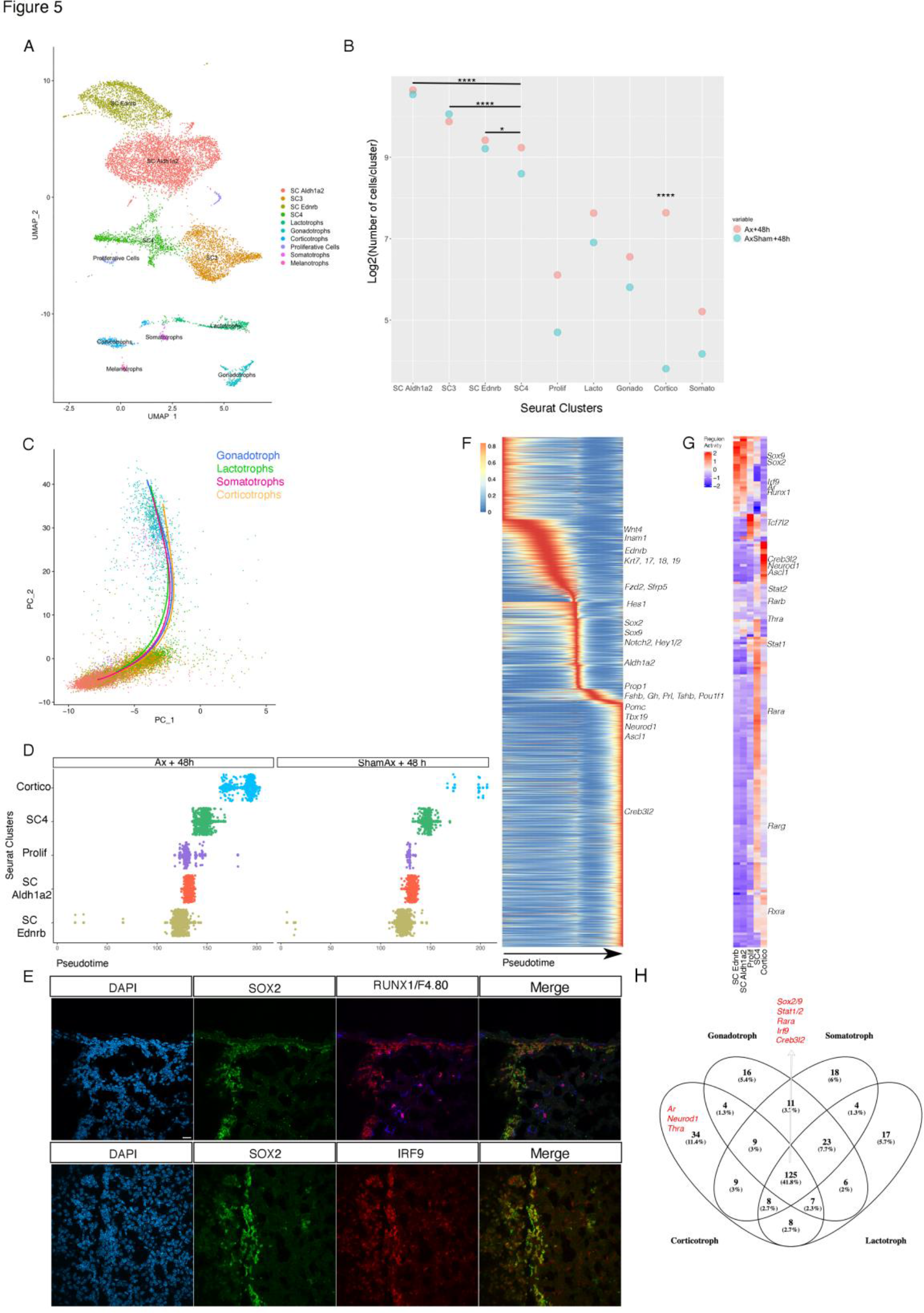
Trajectory analyses following Ax. A) UMAP clustering for integrated datasets from sorted GFP +ve cells from unchallenged, adrenalectomized and sham + 48 h, and + 4 days, all from *Sox9^iresGFP/+^* pituitaries. The data has been filtered to remove mural and vascular cells and outliers. Cluster colour scheme is constant throughout the figure. B) Pairwise comparison for proportion test performed on adrenalectomized and sham + 48h. The proportion of corticotrophs after Ax is significantly different from all other cluster distributions. The proportion of cells in cluster SC4 is significantly different from all other SC cluster distributions (Sup. Table2). C) Representation of endocrine cell trajectories on the PCA plot for integrated dataset. D) Proportion of cells/cluster in the corticotroph trajectory comparing Ax and sham Ax. Differentiation is occurring predominantly after Ax. E) Immunofluorescence on pituitary sections validates RUNX1 and IRF9 expression in SOX2 +ve SCs. RUNX1 is also expressed in F4.80 +ve macrophages. F) Heatmap of 2245 genes associated with the corticotroph trajectory. G) Heatmap of SCENIC analysis on the corticotroph trajectory. H) Venn diagram of the common and unique regulons between the different endocrine trajectories (Sup.Fig. 11).

In sham operated animals, 4.5% of hormone expressing cells, mostly *Prl* +ve, were present, confirming our observations in unchallenged animals (Sup. Table 2). 48 hours after Ax, the proportion of hormone +ve cells had increased to 11%, and this had again doubled after 4 days. Because Sox9iresGFP expression stops as cells differentiate the increasing proportion means that commitment toward differentiation augments during the 4 days following ablation. A pairwise proportion test between the 2 new datasets, sorted identically on the same day, showed that a shift occurs from SC4; while the proportion of cells from each dataset in the 3 SC clusters were comparable, from SC4 the proportion of cells from the Ax dataset becomes significantly increased compared to the control, fitting with its transitioning phase profile (Fig.5B, Sup. Fig.9). At this shorter time point, 48 h00 after adrenal ablation, corticotrophs are the most significantly increased cell type, while 4 days after surgery they were outnumbered by lactotrophs. This sharp and transient increase could represent the specific SC response to adrenals ablation.

We then performed trajectory analyses after principal component analysis (PCA) using Slingshot (*29*) using exclusively Ax+48h and Sham Ax+48h datasets. Trajectories finishing at each endocrine cell type were discovered, with a common root through SC clusters *Ednrb*, *Aldh1a2,* proliferative cells and SC4, then ending at each endocrine cell type (Fig.5C, Sup. Fig.10 and 11). In the corticotroph trajectory (Fig.5D), the expression pattern of 2245 genes significantly correlates with pseudotime (Fig.5F). The expression of known SC markers (*Ednrb*, *Sox2*, *Sox9*, *Aldh1a2*) and signalling pathway members (*Notch2*, *Hes1*, *Hey1* and *Wnt4*, *Fzd2*, *Sfrp5*) was gradually lost. Pathway analyses further suggest keratin filament remodelling, in agreement with epithelial to mesenchymal transition during differentiation. *Ismn1* (*30*) and *Prop1*(*16*), which are involved in cell fate acquisition, are also present during early phases. Conversely, the corticotroph specifier and marker genes *Neurod1*, *Ascl1*, *Tbx19*, *Pomc* and *Creb3l2* (*31*) are upregulated as cells differentiate. A sharp transition occurs as *Prop1* is upregulated and is characterised by a transient up-regulation of *Pou1f1* lineage and gonadotroph (*Fshb*) genes. Comparisons with trajectories from other lineages reveal similar trajectories with a transient up-regulation of unrelated lineages markers (Sup.Fig.11). This could reflect permissive chromatin remodelling as endocrine cell fate acquisition occurs.

SCENIC analyses were then performed to explore GRNs (*32*) and compare them between the different endocrine lineages (Fig.5G, H, Sup.Fig.11). As expected *Sox2* and *9* regulons featured in SC clusters while *Ascl1*, *Neurod1* and *Creb3l2* were present in corticotrophs. The thyroid (*Thra*) and the androgen (*Ar*) hormone receptor regulons featured in corticotroph-differentiating SCs, suggesting that SCs can respond to peripheral signals. New pituitary GNRs were also discovered, such as those involving *Runx1* and *Irf9* whose expression was validated in SCs (Fig.5E, G, H). RUNX1 (RUNt-related transcription tactor 1) is known to control adult skin SC maintenance by regulating WNT signalling (*33*) and is shared between the corticotroph and gonadotroph trajectories. IRFs (Interferon Regulatory Factors) are modulators of cell growth and differentiation beside their anti-viral functions. IRF9 associates with phosphorylated STAT1:STAT2 (Signal Transducer and Activator of Transcription) dimers, and all are present in our four endocrine SCENIC analyses (*34*). Finally, several RA factors were also present in all trajectories, supporting RA involvement during endocrine cell differentiation (Fig.5H).

In conclusion, by adding new datasets, we validated our previous observations suggesting a low differentiation rate of SCs, mostly into lactotrophs in the intact and sham operated animal. We refined the dynamics of the adaptative response after mobilization, showing a transient peak of corticotroph differentiation 48 hours after Ax. Finally, we were able to infer endocrine cell trajectories revealing common and exclusive factors during each endocrine cell fate acquisition.

### Selective maintenance of SC progeny

To examine predictions from our transcriptomic analyses, we performed two different types of lineage tracing experiment. First, we directly assessed expression of hormonal markers on plated *Sox9^iresGFP/+^* FACSorted single cells by immunofluorescence in males and females, 4 days after Ax or sham-surgery. Second, to follow mobilized SCs further, we used *Sox9^ires-CreERT2/+^; R26R^EYFP/+^* animals and examined progeny 10 to 11 days after Cre induction (a week after surgery since induction is mostly performed for 3 days before surgery).

All hormone +ve cell types were found in the GFP +ve fraction in both experimental and control *Sox9^iresGFP/+^* animals, which matches our transcriptomic analyses (Fig.6 A, B). Following Ax, the number of POMC +ve cells increased the most, in agreement with our earliest dataset (Ax+ 48h). However, even if less efficiently stimulated after Ax, the proportion of most other endocrine cell types was comparable to that of corticotrophs. This again reflected our transcriptomic data. More precisely, we also observed a slight increase in somatotroph emergence in both sexes, while lactotrophs and to a reduced extent rare gonadotrophs appear to increase only in females.

**Figure 6:**
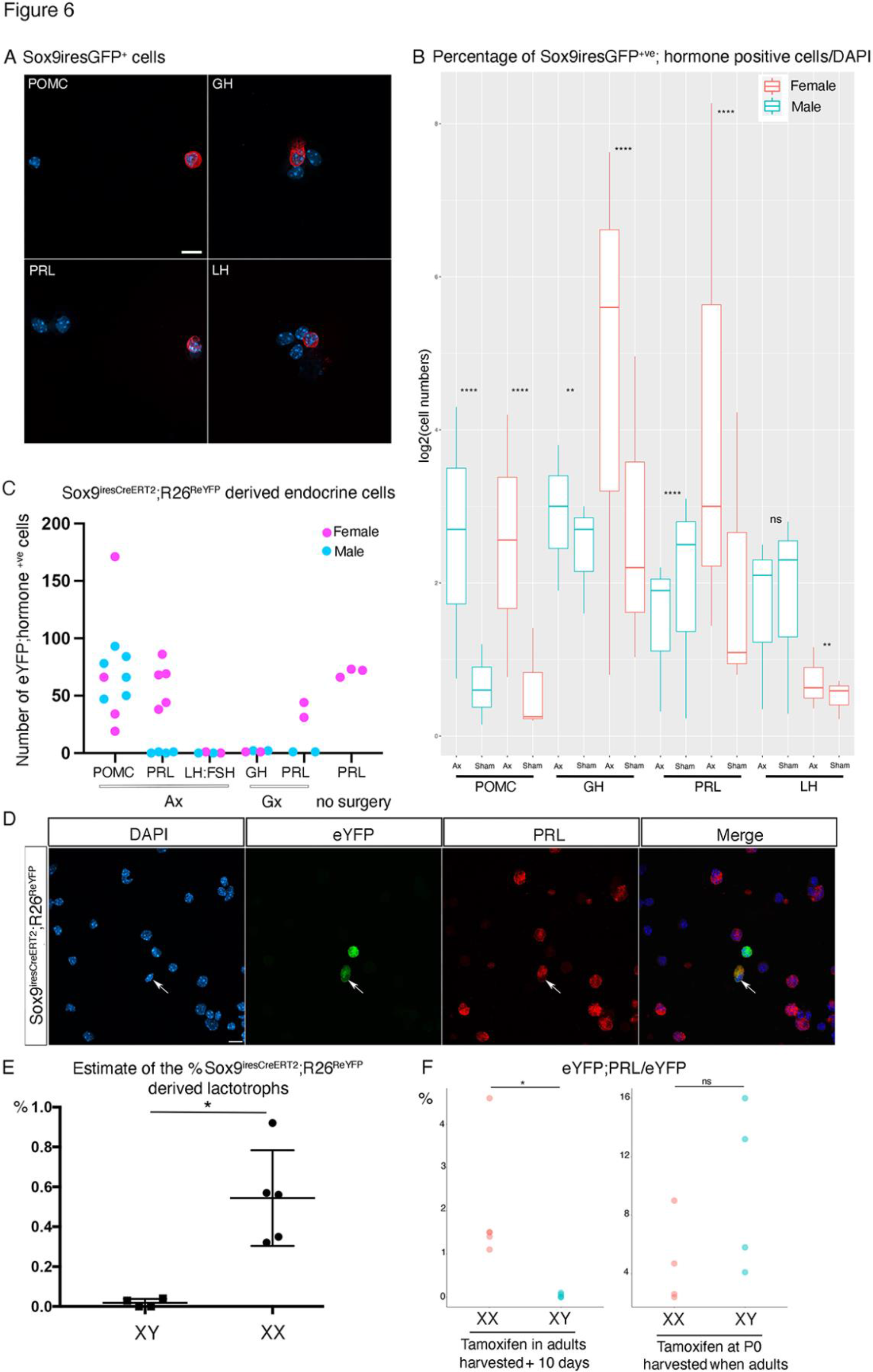
Analyses of scRNAseq predictions by short and long-term lineage tracing. A) Immunofluorescence for hormonal markers on FACsorted Sox9iresGFP +ve cells from a female 4 days after Ax. B) Percentages of FACSorted Sox9iresGFP +ve cells that are hormone +ve /DAPI. Cell counts were performed on plated cells (shown in A). Pituitaries were harvested 4 days after surgeries. Pairwise for proportion tests were performed to assess significance of differences between Ax and Sham (Sup. Table 3). C) Numbers of hormone;eYFP double +ve cells counted in *Sox9^iresCreERT2/+^;Rosa26^ReYFP^*pituitaries a week after surgeries or 10 days after tamoxifen treatment (no surgery). Counts were performed on comparable samples of whole pituitary sections. Corticotrophs are observed in both sexes after Ax, as previously reported. In contrast, significant numbers of lactotrophs are exclusively observed in females. LH;FSH cells have never been observed after gonadectomies (*3*) and were thus not quantified here. D) Immunofluorescence for eYFP and PRL on dissociated non-sorted *Sox9^iresCreERT2/+^;Rosa26^ReYFP^* female pituitary cells. A PRL;eYFP +ve cell is indicated (arrow). E) Estimation of the percentage of lactotrophs differentiated from SCs. eYFP;PRL +ve cells from *Sox9^iresCreERT2^;Rosa26^ReYFP^* dissociated pituitaries were counted 10 days after tamoxifen treatment. Numbers were corrected for Cre induction efficiency. Induction efficiency was calculated in each animal as the percentage of SOX9;eYFP/SOX9 +ve cells present in the cleft, which was preserved from dissociation. There is very little proliferation in normal conditions in the cleft SCs, and we thus assumed that the number of eYFP;SOX9 positive cells directly reflected induction efficiency. 0.6% of lactotrophs originate from SCs 10 days after tamoxifen induction in females while we barely observed any in males (Mann-Whitney test performed on angular transformation of percentages, p=0.0159). F) Percentage of PRL;eYFP/eYFP +ve cells counted 10 days after tamoxifen treatment in 8 to 10 week-old animals, and in 8 week-old animals that were treated with tamoxifen as pups (eYFP;PRL/eYFP in *Sox9^iresCreERT2/+^;Rosa26^ReYFP^*). The proportion of SCs becoming lactotrophs is significantly higher in adult females compared to males (t-test, p=0.03). This sex bias disappears when tamoxifen is administered in pups (t-test test, p=0.1825). Scale bars represent 10 microns.

We then carried out lineage tracing in *Sox9^ires-CreERT2/+^; R26R^EYFP/+^* animals (Fig.6C). We examined the endocrine cell types predicted to be mostly induced by the single cell analysis (Sup.Fig.3C) plus corticotrophs, since we knew these are present in the SC progeny after Ax (*3*) . In contrast with our previous results (Fig.6A), examination of pituitary sections exclusively revealed the presence of POMC;eYFP +ve corticotrophs after Ax, in comparable numbers in both sexes (Fig.6C). In females, we additionally observed lactotrophs, and did so even without performing target organ ablation. Quantification of lactotroph emergence on dissociated pituitaries confirmed the sex bias (Fig.6D). Correcting for CreERT2 induction efficiency, we estimate that in females SCs contribute to 0.6% of the total population of lactotrophs and these SC-derived lactotrophs represent 2% of the eYFP +ve population (Fig.5F).

The two genetic tools we used, *Sox9^iresGFP^* and *Sox9^ires-CreERT2/+^*, display inherent distinctions, such as use of high-dose of the SERM tamoxifen, and purposes, where these may impair direct comparisons. However, while results from both transcriptomic analyses and immunofluorescence demonstrate that endocrine cells of all types are produced in comparable numbers in the short term, we clearly observe that corticotrophs are the only endocrine cells remaining in long-term lineage tracing experiments in males. This indicates a selective maintenance of nascent endocrine cells.

Alternatively, hormone positive cells could acquire expression of SOX9 after surgeries, leading to the presence of SOX9;hormone +ve cells in the SOX9iresGFP fraction but not in the SOX9CreERT2 animals because tamoxifen was given before SC mobilization. There is no supporting evidence for this to happen, while we know that SCs differentiate (SOX9+ve cells become hormone +ve). Furthermore, tamoxifen is known to persist in mice so we should still be able to see these cells in our lineage tracing experiments even if SOX9 becomes up-regulated after the surgeries (*35*). Finally, this discrepancy in the presence of hormone+ve cells in short-term versus long-term lineage tracing already exists in the intact animal because we see these cells our unchallenged and sham single cell RNAseq experiment; but ours and others’ lineage tracing analyses did not reveal any significant differentiation in normal conditions (*3, 4*). Therefore, we think that the most likely explanation is the selective maintenance of a proportion of the SC progeny.

To further explore the sex bias in SC-derived lactotrophs, we administered tamoxifen to P0 *Sox9^ires-CreERT2/+^; R26R^EYFP/+^*pups and quantified differentiation into lactotrophs once they reached adulthood. There was no significant difference in the proportion of SC-derived lactotrophs between males and females (Fig.5E), as previously observed (*3*). Therefore, male SC-derived lactotrophs can be produced, but this is temporally restricted. Furthermore, because the proportion of lactotrophs generated from male and female pups is similar, this suggests that tamoxifen, which can interfere with estrogen action, induces lactotroph emergence. If lactotroph emergence was a continuous process in females their proportion would have been far superior to that observed in males. Our results thus reveal a temporal window of sensitivity in males, while female SCs remain responsive to the lactotroph promoting effect of tamoxifen until adulthood. It is unclear how E2 signalling is altered, and whether SC differentiation is induced or maintenance of nascent lactotrophs permitted. Sexual dimorphism in SC activity has also been reported in another endocrine organ, the adrenals, where androgens regulate cell-turnover (*36*).

### HGF trophic effect on pituispheres suggests that SCs and corticotrophs interact

The selective maintenance of specific nascent SC progeny, while the rest are lost, suggests that survival is dependent on local cues in their microenvironment. In the context of Ax, the corticotrophs are a likely source of signals. We thus re-examined our transcriptomic datasets, screening for signalling pathways potentially activated in SCs by corticotrophs. We found that the receptor tyrosine kinase *Met* is selectively upregulated in SCs while its ligand *Hgf* is present both in corticotrophs and pericytes (Fig.7A-C, Sup.Fig 12 (*37*)), suggesting interactions between these cell types. Furthermore, following Ax, the number of SCs expressing *Met* is increased, exclusively in those proliferating (Fig.7D). This suggests a potential trophic effect of HGF on SCs that we explored in pituisphere assays (Fig.7E, F). Upon HGF treatment, the number of pituispheres is on average doubled compared to the control, demonstrating a trophic effect on SCs. Because the number of corticotrophs is increased after Ax and levels of HGF may thus potentially increase too, this pathway may mediate or at least participate in the increased SC proliferation observed after Ax. In agreement with a trophic role of the pathway in the pituitary, the receptor and its ligand are both widely expressed in human pituitary adenomas (*38*).

**Figure 7:**
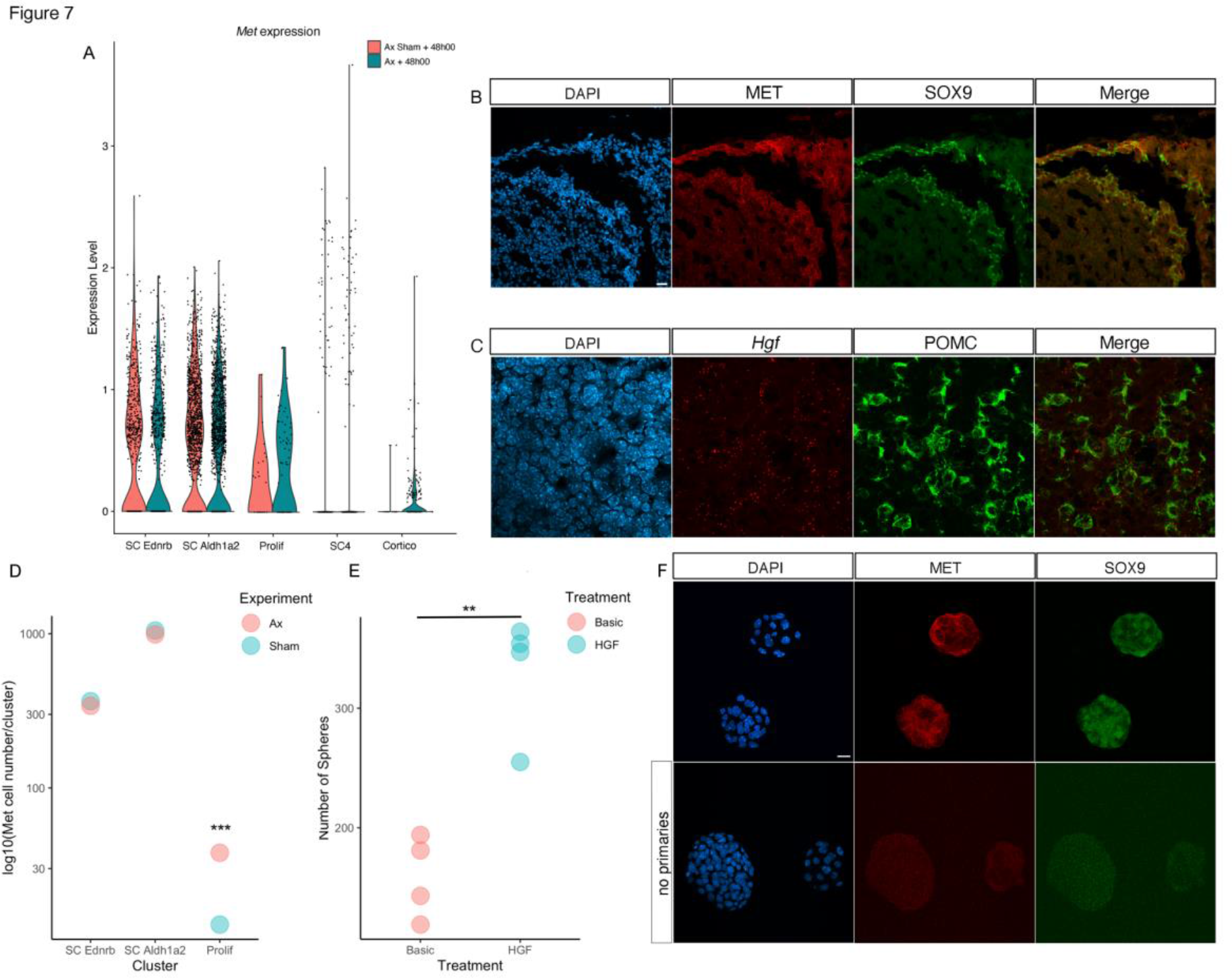
*Met*/MET and *Hgf* are expressed in SCs and corticotrophs respectively and the ligand has a trophic effect on pituispheres. A) Expression levels of *Met* are shown in clusters belonging to the corticotroph trajectory (Fig.5D). B) Immunofluorescence for MET and SOX9 showing the receptor is expressed in SCs. C) In situ hybridisation (RNAscope) and immunofluorescence for *Hgf* and POMC respectively showing that MET ligand is expressed in corticotrophs. *Hgf* is also predicted to be expressed in pericytes which could explain signal outside corticotrophs. D) More cells express *Met* after Ax specifically in the cluster comprising proliferating cells (Chi-squared test, p= 0.0005, Sup.Table 3). E) More pituispheres form when 100ng/ml HGF is added to basic medium (t-test, p= 0.002, Sup. Table 3, n=4). F) MET is expressed in SOX9 +ve pituispheres. Scale bar represents 20 microns in B and D, and 10 microns in C.

## DISCUSSION

While adult pituitary SCs appear superfluous for cell-turnover in the unchallenged animal, their differentiation potential can be stimulated, for example when the function of certain endocrine axes is compromised, such as when adrenals or gonads are ablated (*3*) (*2, 5*). Here we have characterized SC heterogeneity and the modalities of endocrine cell emergence, by examining Sox9iresGFP single cells. We observe that a small percentage of pituitary SCs differentiate in the unchallenged animal, which correlates with single-cell RNAseq analyses performed on both human and mouse pituitaries where the presence of differentiating cells was also detected in the SC compartment (*10*). This proportion increases when SCs are mobilized. The paradigm we applied for SC mobilization, target organ ablation, is known to trigger a specific response in the pituitary: adrenal ablation results in increased ACTH while gonadectomy is associated with increased gonadotropins. However, although there was a significant induction of corticotroph cell fate acquisition shortly after ablation, we unexpectedly observed increased differentiation toward all AL endocrine cell types. Nevertheless, in the longer term, lineage tracing experiments show that it is exclusively the targeted endocrine cell type that is maintained in the SC progeny. Therefore, following differentiation, selective maintenance of specific nascent cells is likely to be an important process regulating pituitary SC endocrine cell output, and it seems likely that this is controlled by the SC microenvironment. Increased sensitivity of nascent pituitary cells to apoptosis had been suggested before (*27*), and shown to increase after target organ ablation, in agreement with our results. Selective fitness of surviving cells in the embryo relies on access to particular trophic factors (*39*). In our Ax paradigm, the pituitary targets of the ablation, the corticotrophs, represent a likely source of signals, a notion re-enforced by the trophic effect of HGF we observe on pituispheres. Similarly, estrogen is known to promote lactotroph proliferation and their emergence in embryos (*40*). We previously examined the effects of estrogen on adult pituitary SC differentiation, but the prior administration of tamoxifen in our lineage tracing system may explain why we did not obtain meaningful results((*3*). Here, our data suggest that tamoxifen treatment induces emergence of lactotrophs, both in male and female neonates, but exclusively in adult females. This in turn implies that this process is sensitive to E2, and that male development suppresses it. The presence of lactotrophs in the male SOX9iresGFP fraction, but not after lineage tracing, suggests that their maintenance is promoted in females. However,to further investigate these aspects a tamoxifen independent lineage tracing tool is required. In parallel with the likely role of corticotrophs on SCs after Ax, lactotrophs may be involved in the selective maintenance of nascent PRL cells emerging after tamoxifen treatment, because they are sensitive to its effect, given that prolactin secretion is increased by the SERM (*41*). In addition to the reliance on trophic factors, it is likely that nascent cell maintenance involves integration into homotypic endocrine networks (*42*). This is reminiscent of what is known for nascent neurons that are often produced in excess and whose persistence relies on synaptic connections (*43*). Although focusing on SCs enabled us to uncover novel regulators and unexpected modalities of the adaptative response, our analyses point toward an important role for the microenvironment, and, in our multi-organ system, peripheric factors which are predicted to modulate the SC response. The thyroid (*Thra*) and the androgen (*Ar*) hormone receptors appear active in differentiating SCs, suggesting that they can respond to peripheral signals, and may integrate this information to regulate pituitary plasticity, as recently proposed (*44*). Further investigations of both local and peripheral interactions are now required to better understand how this adaptative response is regulated in detail.

Differential clustering of SOX9iresGFP +ve SCs, trajectory analyses and *in situ* validation of new markers revealed the activity of region-specific regulatory pathways. In agreement with what had been shown by others (*7*) (*8*), we found two main populations. We show here that these correlate with differential localisation, anterior lobe (ALDH1a2, MSX1 positive) versus more generic AL/IL SC (CLDN3 positive) identity. This suggests that AL and IL SCs are regulated differently. We speculate that RA may be involved in AL SC differentiation, as shown in the embryo (*11*). The downregulation of ALDH1a2 expression in IL SCs which do not differentiate (*2, 45*) provides an additional support to this hypothesis. The enrichment of EDNRb in the cells lining the cleft reveals a further level of heterogeneity between cleft lining and parenchymal cells. In the trajectory analyses, the upstream expression pattern of *Ednrb* implies that the SCs lining the cleft are in a more immature state than those in the parenchyma, which fits with this epithelium constituting the primary pituitary SC niche. Among the 3 endothelin ligands, *Edn3* is the most abundantly expressed in the gland, more precisely in SCs and pericytes (*37*). This suggests that this pathway could function in an autocrine or paracrine way in cleft SCs, because these are not in close contact with capillaries (*42*), which would make an interaction with the latter unlikely. EDNRB signalling has been shown to regulate melanocyte SC regeneration (*46*) and we are currently investigating whether this role is conserved in pituitary SCs. We have furthermore found that a subpopulation of parenchymal SCs is defined by AQP3 expression. It is unclear whether this correlates with the existence of functionally different subpopulations, such as supporting FS cells versus SCs for example, or whether this reflects distinct SC states, but we are now able to identify these cells, and investigate their role. Finally, the presence of RUNX, IFR and STAT regulons during cell differentiation is also of potential interest because of the roles of these factors in different SC contexts, and because they have been shown to interact (RUNX/STAT and IFR/STAT). STAT transcription factors are activated by different cytokines and growth factors, in particular IL6 whose role in the regenerative response of pituitary SCs was recently uncovered (*7*).

Our differentiation trajectories suggest the presence of a common root for all endocrine cell types, which only segregate as the terminal differentiation markers are upregulated in the nascent cells. This process is likely to be fast since cells still contain GFP expressed from *Sox9iresGFP*. It is now important to determine whether the same genes in this common root are associated with the different lineages and when specificity first appears. It is tempting to draw a parallel with embryonic pituitary progenitors, because these were similarly shown to all commit at the same stages (*47*). Interestingly, it was proposed that intrinsic factors and cell communication within the developing pituitary underlie specific endocrine cell fate acquisition, which, in a different context and with different modalities and outcomes, is what we hypothesize is happening in our model.

In conclusion, evaluation of single cell transcriptomic predictions with short and long-term lineage tracing experiments has proved a robust strategy revealing novel aspects of pituitary SC adaptative responses. In circumstances where an exceptional pathophysiological need is perceived, nascent cells of the appropriate type are maintained. Understanding how the new cells can survive will help us to understand how SC input is controlled in the pituitary, but this could also be relevant in other tissues where SC activity appears largely superfluous. Finally, it is notoriously difficult to obtain mature cell types *in vitro* for use in regenerative medicine. Therefore, identifying trophic factors for nascent cells, which may be all that can be obtained for transplantation, appears necessary.

## Supporting information

All supplementary figures and tables

## Acknowledgements

We are grateful for the support from present and past members of the Lovell-Badge lab. We are indebted to the Francis Crick Institute platforms for their excellent support and expertise. We especially thank Jacek Mor from the Biological Research facility for taking excellent care of the animals, Donald Bell from the Advanced Light microscopy, Deborah Schneider-Luftman from Bioinformatics and Biostatistics and experts from the Advanced Sequencing and Histology platforms. We also thank Frederique Ruf-Zamojski and Zidong Zhang from the Icahn School of Medicine at Mount Sinai (New York) for sharing their expertise. This work was supported by the Medical Research Council, U.K. (U117512772, U117562207 and U117570590) and the Francis Crick Institute which receives its core funding from Cancer Research UK (FC001107), the UK Medical Research Council (FC001107), and the Wellcome Trust (FC001107).

## SUPPLEMENTARY FIGURE LEGENDS

<u>Sup. Figure 1: Expression of known and novel markers in SOX9iresGFP unchallenged dataset.</u>

A) *Sox2* and *Sox9* were used to identify SCs while genes encoding for hormones (*Gh*, *Prl*, *Tshβ*, *Lhβ*, *Fshβ*, *Pomc*) and lineage specific transcription factors were examined to distinguish committed cells (*Pou1f1* for somatotrophs, lactotrophs and thyrotrophs, *Tbx19* and *Pax7* for corticotrophs and melanotrophs, and *Nr5a1* for gonadotrophs). B) Heatmap for the top10cluster markers.

<u>Sup. Figure 2: Co-localisation between SOX9 and ACTH after Ax.</u>

Co-immunofluorescence for GFP, SOX9 and ACTH on sections of pituitaries harvested one week after Ax. Expression of Sox9iresGFP (A) and SOX9 (B) is observed in rare ACTH +ve cells (arrows) exclusively after Ax. The scale bar represents 10 μm for A and 5μm for B.

<u>Sup. Figure 3: Gating strategy for SOX9iresGFP cells.</u>

Gating for SOX9iresGFP cells was validated using wild-type animals. The SOX9iresGFP fraction we selected typically represented 1 to 2% of all cells while 0.02% of all cells were falsely sorted in negative controls. This contamination of likely hormonal cells represents a small proportion of the hormonal cells we detect in our unchallenged Sox9iresGFP fraction (3% of 1.9% representing 0.08% of the whole population).

<u>Sup. Figure 4: Expression of novel SC cell markers in whole pituitary and SC datasets.</u>

<u>A) UMAP of re-analysed whole pituitary dataset</u> (*28*). This dataset was used to examine levels of expression of *Ednrb* (B), *Aldah1a2* (C), *Cldn3* (D) and *Msx*1(E) to show exclusive or enriched expression in SCs. F) Co-immunofluorescence for MSX1, GFP and SOX9 in a male SOX9iresGFP pituitary section. MSX1 is exclusively expressed in SOX2;SOX9 AL SC (yellow arrow shows its absence from IL SC, and white arrow shows its expression in parenchymal SC). *Msx1* violin plot shows enrichment in cluster 0 of the single cell dataset presented in Fig.1. The scale bar in A represents 10 μm.

<u>Sup. Figure 5: In situ hybridisation for *Ednrb*.</u>

In situ hybridisation analysis (RNAscope) of *Ednrb* expression confirming higher levels of expression in cleft versus parenchymal SC where low levels of signal can be detected, while levels of *Sox9* expression appear similar in both compartments. The scale bar represents 200 μm.

<u>Sup. Figure 6: Expression of FS cells markers in SC and whole pituitary datasets.</u>

A-C) UMAP for *S100b* (A), *Aldolase C* (B) and *Slc15a2* (C) in our Sox9iresGFP single cell dataset.

D-F) UMAP for *S100b* (D), *Sox2* (E) and *Aldolase C* (F) in a whole pituitary dataset (*28*).

<u>Sup. Figure 7: Unchallenged/Challenged integrated analysis.</u>

A) UMAP clustering and marker analysis for integrated datasets from sorted *Sox9^iresGFP/+^* cells from unchallenged, adrenalectomized, gonadectomized and both adrenalectomized and gonadectomized animals, 4 days after surgery.

B) Pairwise comparison for proportion test (‘pairwise_prop_test’ function [stats R package]) on stem cell clusters (circled on UMAP in A). The number of cells assigned to cluster 10 corresponding to proliferative cells (see *Top2a* expression in A) is significantly higher in activated stem cells (Sup. Table2).

C) Numbers and proportion of specific hormone +ve cells/all hormone +ve cells in each dataset. While we would expect the number of hormone +ve cells to increase after target organ ablation, as seen here, the fact that we select fewer GFP weak cells in unchallenged animals prevents us from comparing the proportion of hormone +ve cells in unchallenged and mobilized datasets and thus make this conclusion.

<u>Sup. Figure 8: UMAP clustering for filtered integrated analysis split by dataset.</u>

A) UMAP clustering for integrated datasets from sorted SOX9iresGFP +ve cells from unchallenged, adrenalectomized and sham + 48 h, adrenalectomized + 4 days split by dataset. SC clusters 3 and 4 mostly originate from datasets collected 48 hours after surgery.

<u>Sup. Figure 9: Analyses of marker expression on UMAP clustering in Ax integrated datasets.</u>

Markers of SC and hormone-secreting cell types are shown on the UMAP clustering for integrated datasets from GFP +ve cells sorted from *Sox9^iresGFP/+^* pituitaries from mice that were unchallenged, adrenalectomized and sham + 48 h, adrenalectomized + 4 days.

<u>Sup. Figure 10: PCA clustering for filtered integrated analysis split by dataset.</u>

SC4 cluster cells located in an intermediate position between SC and endocrine cells originate mostly from Ax+48h, which is consistent with mobilization and increased commitment toward differentiation.

<u>Sup.Fig.11: Analysis of endocrine trajectories and regulons.</u>

Heatmaps of genes associated with trajectories and SCENIC analyses for lactotroph, somatotroph and gonadotroph lineages. Representative genes are shown. Known regulators and those in common with the corticotroph analyses (Fig.5 F-H) are displayed on the heatmaps.

<u>Sup. Figure 12: Expression of *Met* and *Hgf* in the whole pituitary dataset.</u>

Expression of *Hgf* was plotted along with *Pomc* and *Pdgfrb* suggesting expression in *Pomc* positive corticotrophs and *Pdgfrb* positive pericytes, while *Met* expression is restricted to *Sox2* positive SCs (from re-analysed dataset (*28*)). The same pattern is observed in a different dataset (*37*).

<u>Sup. Table1: Pathway analysis for clusters 0 and 2.</u>

Gene markers for WT Cluster 0 were used to determine enriched pathways with Metacore using a hypergeometric test. Pathways with an adjusted p value < 0.05 are shown in the table. Genes present in our dataset are highlighted in the top5 pathways.

<u>Sup. Table2: Endocrine cell cluster numbers from Ax integrated dataset (Fig.5)</u>

<u>Sup. Table 3: Pairwise for proportion test results.</u>

<u>Sup. Table 4: Counting numbers for Fig. 2, 3, 5 and 6.</u>

<u>Sup. Table 5: Cell type signatures.</u>

## MATERIALS AND METHODS

### Mice

All animal experiments carried out were approved under the UK Animals (Scientific Procedures) Act 1986 and under the project licences n. 80/2405 and PP8826065. *Sox9^iresGFP/+^* (*Sox9^tm1Haak^*) (*48*), *Sox9^iresCreERT2/+^* (*Sox9^tm1(cre/ERT2)Haak)^* (*48*), *Wnt1Cre ((no gene)^tg(Wnt1-GAL4)11Rth^*) (*49*) and Rosa26^ReYFP/ReYFP^ (*Gt(ROSA)26Sortm1(EYFP)Cos*) (*50*) mice were maintained on a C57Bl6 background.

For CreERT2 induction in *Sox9^iresCreERT2/+^* ;Rosa26^ReYFP/+^ animals, tamoxifen was administered for 3 consecutive days at a concentration of 0.2mg/g body weight. For *Sox9^iresCreERT2/+^* ;Rosa26^ReYFP/+^ pups induction, a single subcutaneous injection of tamoxifen (0.25mg/g body weight) was performed. Target organ ablations were performed as previously described (*3*).

### Pituitary SC selection

The anterior lobe of freshly dissected 8 to 12 week-old pituitaries was minced and incubated for 15mn at 37°C in 1mg/ml Papain (Sigma 10108014001), 10μg/ml DNAse I (Sigma 10104159001) and 10μg/ml Rock inhibitor (Abmole Bioscience M1817) in HBSS (Sigma H9394). pH was adjusted by addition of NaOH. The enzymatic solution was removed, and the cells were mechanically resuspended in HBSS supplemented with DNAse I and Rock inhibitor, as above. Cells were FACsorted for GFP using a large nozzle (100μm) and low pressure (20psi) for optimal cell survival. For immunofluorescence, sorted cells were plated on Superfrost slides for 90 min in a cell culture incubator, fixed 20 min on ice in 4% PFA and immediately processed.

### Pituisphere culture

Dissociated anterior lobes (see above) of male and female 7 to 12 week-old mice were seeded at 50.10^3^ cells/ml in pituisphere medium containing EGF and FGF (*9*). 100ng/ml of HGF (2207-HG-025/CF, R&D systems) was added to this basic medium in treated cultures.

Medium was replaced every other day and spheres manually and blindly counted after 7 days. 4 independent repeats were performed.

### Single cell sequencing

The concentration and viability of the single cell suspension was measured using Trypan Blue and the Eve automatic cell counter (unchallenged and target organ ablation d4 samples) or acridine orange and propidium iodide with the Luna-FX7 automatic cell counter (Ax and Sham control d2 samples). Up to 10,000 cells were loaded on Chromium Chip and partitioned in nanolitre scale droplets using the Chromium Controller and Chromium 10x 3 prime v2.0, Chromium Single Cell 3’ Reagent Kits User Guide (v2 Chemistry) (unchallenged and target organ ablation d4 samples) or Next GEM Single Cell reagents (3 prime v3.1) (Ax and Sham control d2 samples). Within each droplet the cells were lysed, and the RNA was reverse transcribed. All of the resulting cDNA within a droplet shared the same cell barcode. Illumina compatible libraries were generated from the cDNA using Chromium Next GEM Single Cell library reagents in accordance with the manufacturer instructions (10x Genomics). Final libraries are quality checked using the Agilent TapeStation and sequenced using the Illumina HiSeq4000 or HiSeq2500, read configuration 26-8-0-98 (unchallenged and target organ ablation d4 samples) or NovaSeq 6000, read configuration 28-10-10-90. Where expression of specific genes in the whole pituitary gland is referenced as ((*37*), we used this published dataset on the Loupe browser to examine expression of relevant markers.

### Bioinformatic analyses

Alignment, Seurat analysis, integration and cluster annotation were performed using CellRanger and Seurat. Raw reads were initially processed by the Cell Ranger v.3.0.2 pipeline (*51*), which deconvolved reads to their cell of origin using the UMI tags, aligned these to the mm10 transcriptome (to which we added the eGFP sequence https://www.addgene.org/browse/sequence/305137/ to detect eGFP expressing cells) using STAR (v.2.5.1b) (*52*) and reported cell-specific gene expression count estimates. All subsequent analyses were performed in R v.3.6.0 (*53*) using the Seurat (v3) package (*54*). Genes were considered to be ‘expressed’ if the estimated (log10) count was at least 0.1. Primary filtering was then performed by removing from consideration cells expressing fewer than 50 genes and cells for which mitochondrial genes made up greater than 3 standard deviations from the mean of mitochondrial expressed genes. PCA decomposition was performed and, after consideration of the eigenvalue ‘elbow-plots’, the first 20 components were used to construct the UMAP plots per sample. Multiple samples, generated in this study and published (*28*), were integrated using 2000 variable genes and Seurat’s CCA method. Cluster specific gene markers were identified using a Wilcoxon rank sum test and the top 20 genes ranked by logFC per cluster were used to generate a heatmap. Clusters were annotated using label tranfer methods within Seurat using the GSE120410 dataset (*28*). Clusters were further annotated using cell specific signatures (Sup. Table 5).

Cell trajectories were identified using the package ’Slingshot’ (version 1.4.0) (*29*), using the undifferentiated cluster as a starting point and the PCA co-ordinates. Lineages were identified showing specific trajectories ending in specific differentiated cells. Differential expression of genes between samples were further investigated for the corticotroph lineage with a correlation metric against pseudotime, using the ’gam’ package (version 1.20) (*55*). Heatmaps were made using the smoothed expression of differential genes, and the cells are ranked by increasing pseudotime.

In order to investigate transcription factor activity along pseudotime, the package SCENIC (*32*) was used. A list of 948 putative mouse regulatory binding sites found in promoter regions of expressed genes were used to identify shared regulatory networks. Regulon activity score per transcription factor was calculated using the AUCell function, where the enrichment of target genes was measured using the area under the curve of the target gene relative to the expression-based ranking of all genes. The top 200 regulons ranked by activity score were using to generate a binarized heatmap, where the cells were ranked by pseudotime.

### Immunofluorescence and RNAscope

Pituitaries were fixed after dissection by immersion in 4% PFA at 4°C. Immunofluorescent stainings were performed on cryosections as described (*56*). The following primary antibodies were used: Goat anti-SOX2 (Biotechne, AF2018), rabbit anti-SOX9 (a gift from F. Poulat, IGH Montpellier, France), rat anti-GFP (Nacalai Tesque 04404-84), chick anti-GFP (Invitrogen, A10262), rabbit anti-EDNRb (Abcam 117529), goat anti-MSX1 (R&D system AF5045), rabbit anti-ALDH1A2 (Abcam 75674, after antigen retrieval), rabbit anti-Claudin3 (Abcam ab15102), rat anti-PECAM (BD Pharmigen, 550274), rat anti-PDGFRβ (eBiosciences 14-1402-81), rabbit anti-AQP3 (Alomone Labs, AQP-003, after antigen retrieval) rabbit anti-Runx1 (Sigma, HPA004176), rabbit anti-IRF9 (Cell Signalling, D9I5H), goat anti-Met (R&D, AF 527) and hormone antibodies anti-LH, GH, ACTH and PRL from the National Hormone Peptide Program (A.F. Parlow, Torrance, USA). Slides were then incubated with Alexa Fluor secondary antibodies. Imaging of stained tissue sections was performed on a Leica SPE microscope while imaging of plated cells was done on an Olympus spinning disk microscope.

RNAscope was performed on cryosections following the manufacturer’s instructions using *Hgf* (#456511) and *Ednrb* (#473801) probes.

### Cell counts

Counts were performed blindly. For dissociated cell counts, slides stained by immunofluorescence were scanned; automated counts were then performed using QuPath (https://qupath.github.io/). For *Sox9^iresCreERT2/+^* ;Rosa26^ReYFP/+^;hormone counting on sections, double +ve cells were manually identified and each imaged on a slide representing 1/5 of a pituitary. Images were then reviewed, and double +ve cells counted. All counting results and tests are presented (Sup. Table 3 and 4).

### Statistics

Comparisons of the percentages of Sox9;PDGFRβ (Fig.3) were performed after angular transformation using Mann-Whitney test on Prism. Pairwise for proportion test (Sup. Fig4B, Fig.5B, Fig.6B and Sup. Table 3), t-test (Fig.6E, Fig.7D) and Chi squared test (Fig.7) were performed using the Stats R package.

## Author contributions

Conceptualization: KR, RLB

Methodology: KR

Bioinformatic analyses: PC

Investigation: KR, DS

Visualization: KR

Supervision: KR, RLB

Writing—original draft: KR

Writing—review & editing: KR, RLB

## Competing interests

Authors declare that they have no competing interests.

## Data and materials availability

All data are available in the main text or the supplementary materials.

Codes are available on request. Datasets have been submitted (Awaiting confirmation of data deposited at GEO).

## Notes

### Competing Interest Statement

The authors have declared no competing interest.

### Summary of Updates

We first ascertained that the differentiated cells we analysed were indeed present in the Sox9iresGFP positive fraction. This was done through the use of additional controls, and we present this new data. Because some of our conclusions were based on the presence of small percentages of differentiated cells, it was important to rule out the possibility of contamination. Secondly, we have analysed additional markers and found a subpopulation of AQP3 positive parenchyma stem cells, revealing further heterogeneity in the SOX9;SOX2 positive stem cell population. Thirdly, we have found that the MET/HGF pathway, which we previously hypothesized may regulate the SC trophic response, induces the formation of an increased number of pituispheres in vitro, in agreement with this hypothesis. This strongly supports our main proposition of an instructive role of the pituitary microenvironment for stem cell mobilization. Furthermore, we have defined cell trajectories and regulons for all the endocrine lineages and compared these to identify common and specific regulators.

