## Supplementary material for "SOX9-positive pituitary stem cells differ according to their position in the gland and maintenance of their progeny depends on context": All supplementary figures and tables

Sup.Fig.1

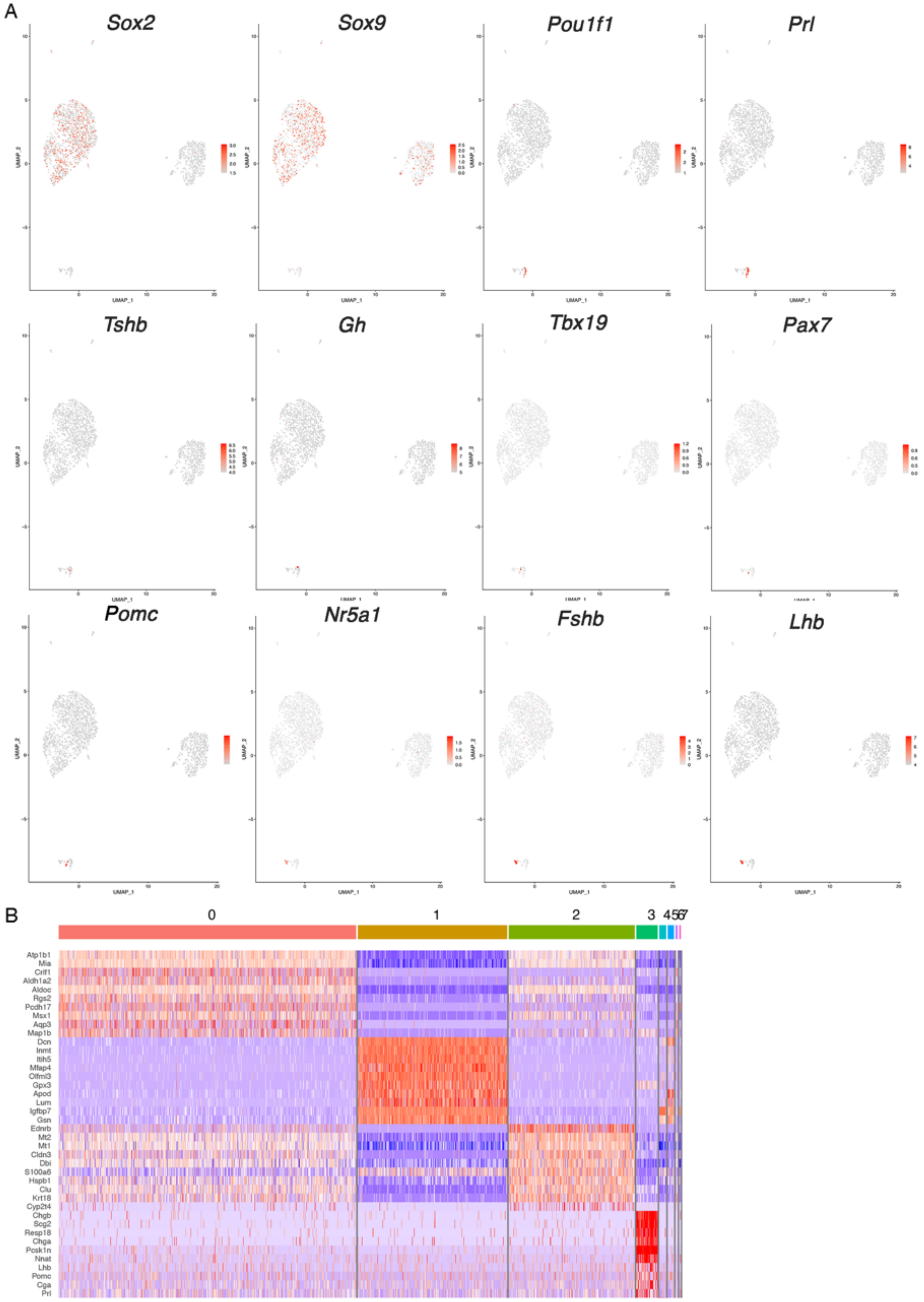
A
Sox2
Sox9
Pou1f1
Prl
Tshb
Gh
Tbx19
Pax7
Pomc
Nr5a1
Fshb
Lhb
UMAP_1
UMAP_2
B
0
1
2
3
4567
Atp1b1
Mia
Crlf1
Aldh1a2
Aldoc
Rgs2
Pcdh17
Msx1
Aqp3
Map1b
Dcn
Inmt
Itih5
Mfap4
Olfml3
Gpx3
Apod
Lum
Igfbp7
Gsn
Ednrb
Mt2
Mt1
Cldn3
Dbi
S100a6
Hspb1
Clu
Krt18
Chgb
Scg2
Resp18
Chga
Pcsk1n
Nnat
Lhb
Pomc
Cga
Prl

Fig. S1. Expression of known and novel markers in SOX9iresGFP unchallenged dataset.

A) *Sox2* and *Sox9* were used to identify SCs while genes encoding for hormones (*Gh*, *Prl*, *Tshβ*, *Lhβ*, *Fshβ*, *Pomc*) and lineage specific transcription factors were examined to distinguish committed cells (*Pou1f1* for somatotrophs, lactotrophs and thyrotrophs, *Tbx19* and *Pax7* for corticotrophs and melanotrophs, and *Nr5a1* for gonadotrophs). B) Heatmap for the top10cluster markers.

Sup. Fig.2

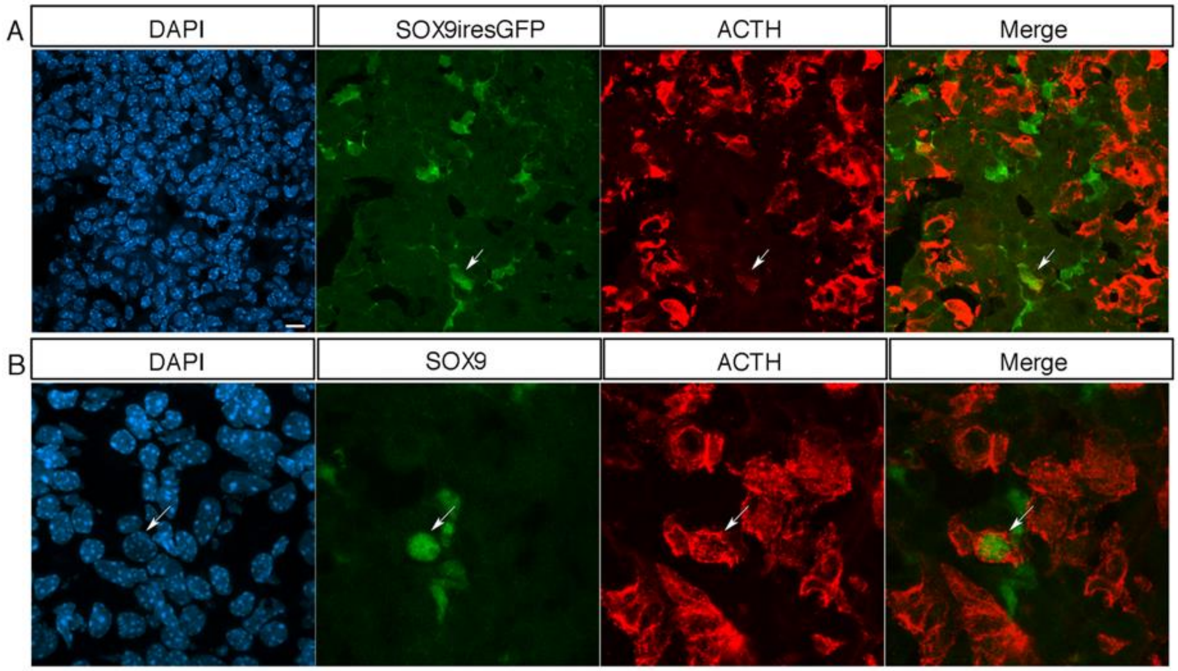

Fig. S2. <u>Co-localisation between SOX9 and ACTH after Ax.</u>

Co-immunofluorescence for GFP, SOX9 and ACTH on sections of pituitaries harvested one week after Ax. Expression of Sox9iresGFP (A) and SOX9 (B) is observed in rare ACTH +ve cells (arrows) exclusively after Ax. The scale bar represents 10 μm for A and 5μm for B.

Sup. Fig.3

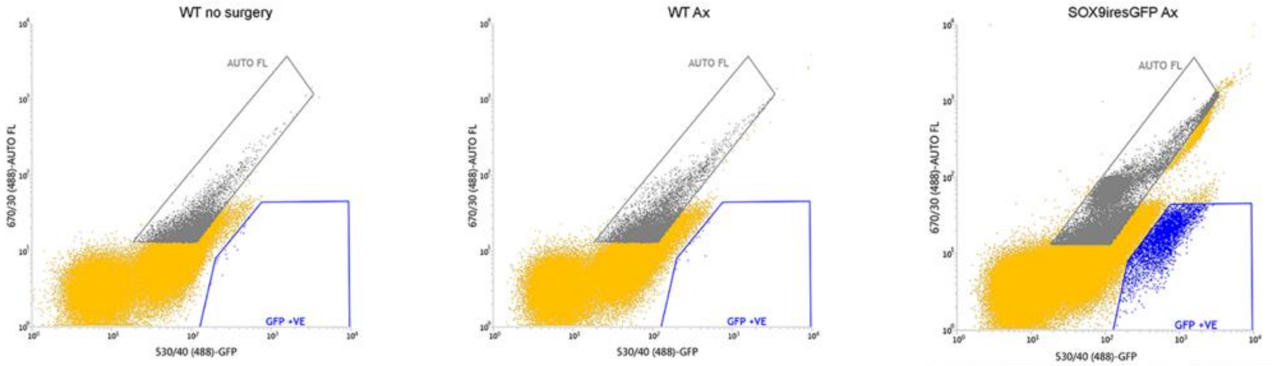

Statistics: 20230126 KR INFLUX A_CONTROL

| Populations | Events | % Total | % Parent | FSC Mean | SSC Mean |
|---|---|---|---|---|---|
| All Events | 124,738 | 100.00% | #### | 14,605 | 11,663 |
| P1 SCATTER GATE | 102,436 | 82.12% | 82.12% | 16,890 | 13,578 |
| P2 SINGLET GATE | 100,000 | 80.17% | 97.62% | 16,840 | 13,448 |
| GFP +VE | 24 | 0.02% | 0.02% | 7,634 | 9,554 |
| AUTO FL | 12,098 | 9.70% | 9.70% | 20,590 | 19,283 |

Statistics: 20230126 KR INFLUX A_SAMPLE Ax

| Populations | Events | % Total | % Parent | FSC Mean | SSC Mean |
|---|---|---|---|---|---|
| All Events | 132,584 | 100.00% | #### | 13,028 | 9,199 |
| P1 SCATTER GATE | 102,041 | 76.96% | 76.96% | 15,850 | 11,445 |
| P2 SINGLET GATE | 100,000 | 75.42% | 98.00% | 15,846 | 11,393 |
| GFP +VE | 23 | 0.02% | 0.02% | 22,527 | 22,377 |
| AUTO FL | 11,526 | 8.69% | 8.69% | 18,968 | 15,292 |

Statistics: 20230126 KR INFLUX A_GFP SAMPLE

| Populations | Events | % Total | % Parent | FSC Mean | SSC Mean |
|---|---|---|---|---|---|
| All Events | 263,026 | 100.00% | #### | 13,998 | 10,498 |
| P1 SCATTER GATE | 195,761 | 74.43% | 74.43% | 17,485 | 13,472 |
| P2 SINGLET GATE | 192,598 | 73.22% | 98.38% | 17,436 | 13,447 |
| GFP +VE | 5,000 | 1.90% | 2.60% | 14,537 | 3,804 |
| AUTO FL | 38,839 | 14.77% | 14.77% | 19,133 | 17,443 |

Fig. S3. Gating strategy for SOX9iresGFP cells.

Gating for SOX9iresGFP cells was validated using wild-type animals. The SOX9iresGFP fraction we selected typically represented 1 to 2% of all cells while 0.02% of all cells were falsely sorted in negative controls. This contamination of likely hormonal cells represents a small proportion of the hormonal cells we detect in our unchallenged Sox9iresGFP fraction (3% of 1.9% representing 0.08% of the whole population).

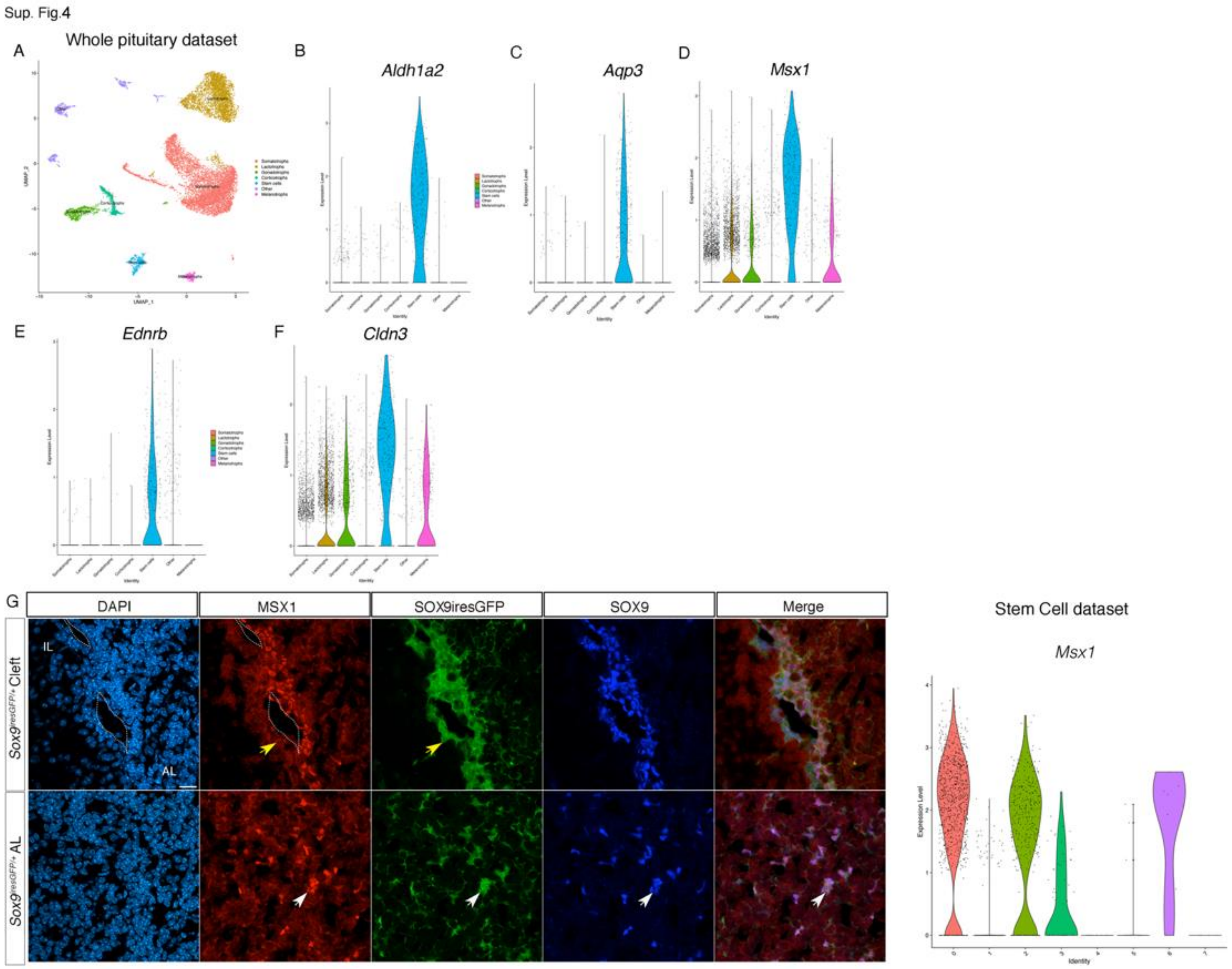

Fig. S4. Expression of novel SC cell markers in whole pituitary and SC datasets.

A) UMAP of re-analysed whole pituitary dataset (*28*). This dataset was used to examine levels of expression of *Ednrb* (B), *Aldah1a2* (C), *Cldn3* (D) and *Msx*1(E) to show exclusive or enriched expression in SCs. F) Co-immunofluorescence for MSX1, GFP and SOX9 in a male SOX9iresGFP pituitary section. MSX1 is exclusively expressed in SOX2;SOX9 AL SC (yellow arrow shows its absence from IL SC, and white arrow shows its expression in parenchymal SC). *Msx1* violin plot shows enrichment in cluster 0 of the single cell dataset presented in Fig.1. The scale bar in A represents 10 μm.

Sup.Fig.5

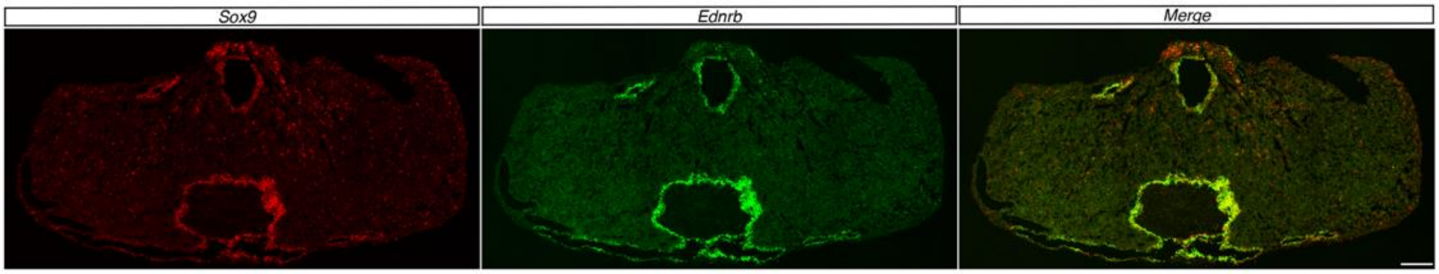

Fig. S5. <u>In situ hybridisation for *Ednrb*.</u>

In situ hybridisation analysis (RNAscope) of *Ednrb* expression confirming higher levels of expression in cleft versus parenchymal SC where low levels of signal can be detected, while levels of *Sox9* expression appear similar in both compartments. The scale bar represents 200 μm.

Sup. Fig.6

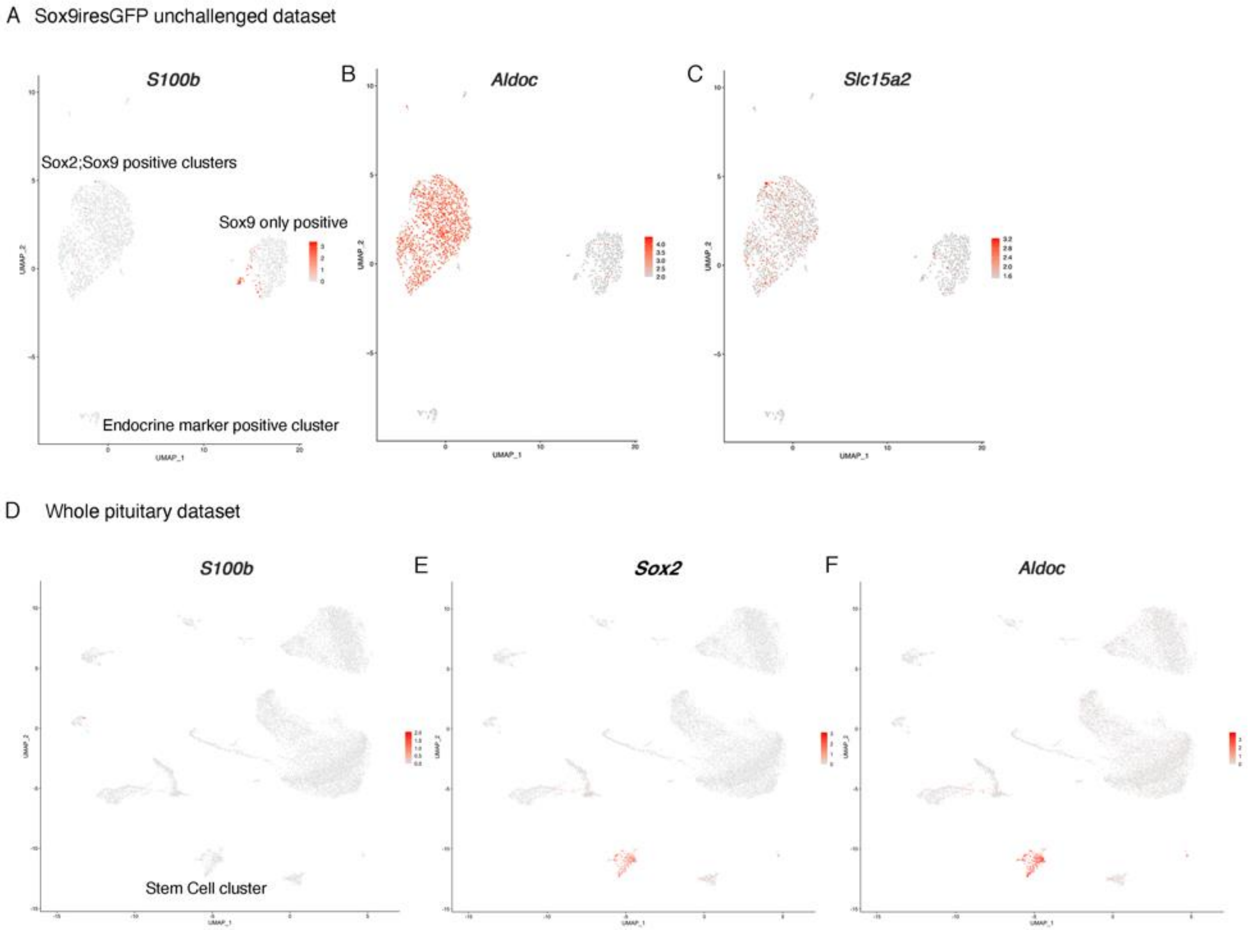

Fig. S6. Expression of FS cells markers in SC and whole pituitary datasets.

A-C) UMAP for *S100b* (A), *Aldolase C* (B) and *Slc15a2* (C) in our Sox9iresGFP single cell dataset.
D-F) UMAP for *S100b* (D), *Sox2* (E) and *Aldolase C* (F) in a whole pituitary dataset (*28*).

Sup. Fig.7

A

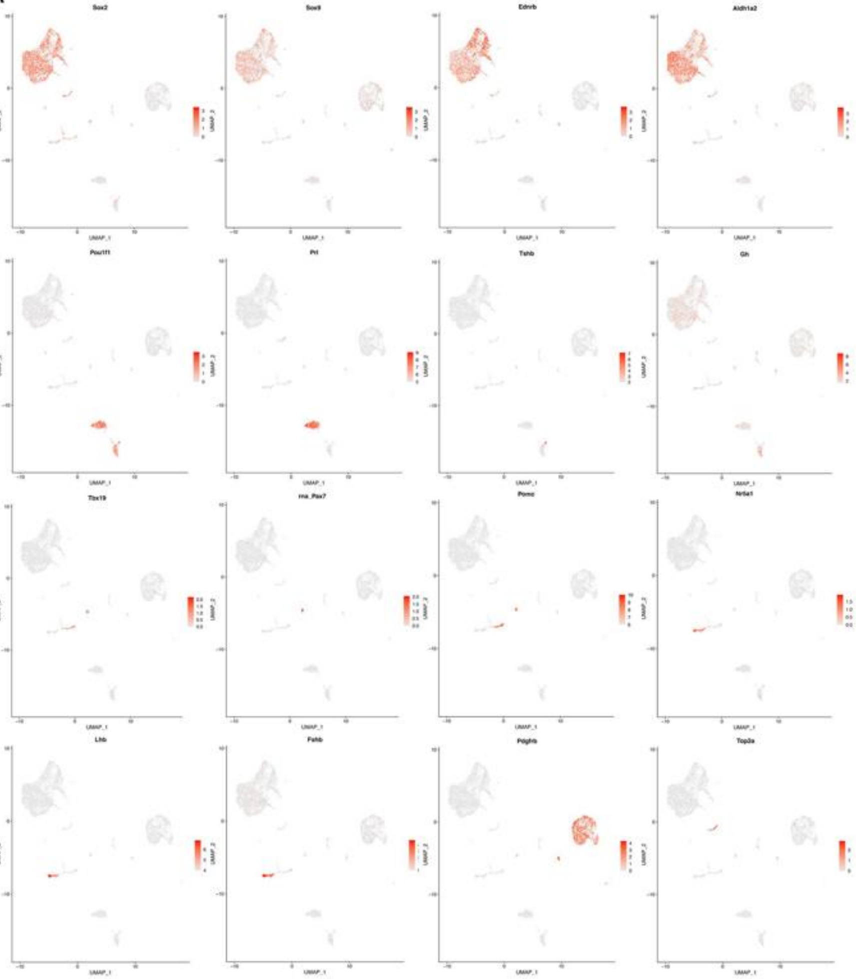
Sox2
Sox9
Ednrb
Aldh1a2
Pou1f1
Prl
Tshb
Gh
Tbx19
Pomc
Nr5a1
Lhb
Fshb
Top2a
UMAP_1
UMAP_2

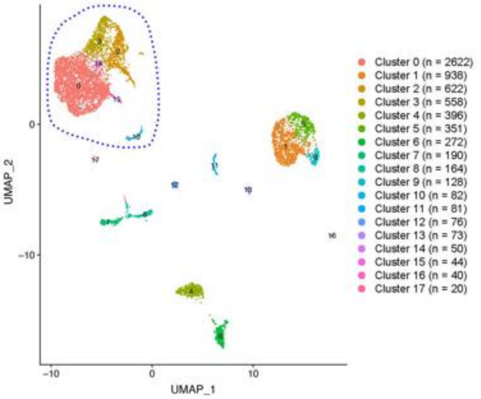
UMAP_1
UMAP_2
Cluster 0 (n = 2622)
Cluster 1 (n = 938)
Cluster 2 (n = 622)
Cluster 3 (n = 558)
Cluster 4 (n = 396)
Cluster 5 (n = 351)
Cluster 6 (n = 272)
Cluster 7 (n = 190)
Cluster 8 (n = 164)
Cluster 9 (n = 128)
Cluster 10 (n = 82)
Cluster 11 (n = 81)
Cluster 12 (n = 76)
Cluster 13 (n = 73)
Cluster 14 (n = 50)
Cluster 15 (n = 44)
Cluster 16 (n = 40)
Cluster 17 (n = 20)

B

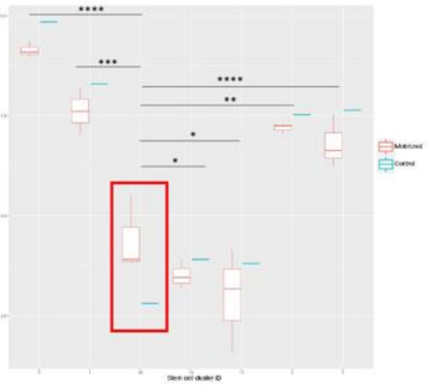
****
***
****
**
*
*
Mobilized
Control
Stem cell cluster ID

C

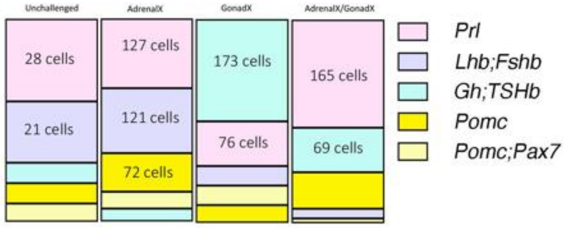
Unchallenged
AdrenalX
GonadX
AdrenalX/GonadX
28 cells
21 cells
127 cells
121 cells
72 cells
173 cells
76 cells
165 cells
69 cells
Prl
Lhb;Fshb
Gh;TSHb
Pomc
Pomc;Pax7

Fig. S7. Unchallenged/Challenged integrated analysis.

A) UMAP clustering and marker analysis for integrated datasets from sorted *Sox9*$^{iresGFP/+}$ cells from unchallenged, adrenalectomized, gonadectomized and both adrenalectomized and gonadectomized animals, 4 days after surgery.
B) Pairwise comparison for proportion test ('pairwise_prop_test' function [stats R package]) on stem cell clusters (circled on UMAP in A). The number of cells assigned to cluster 10 corresponding to proliferative cells (see *Top2a* expression in A) is significantly higher in activated stem cells (Sup. Table2).

Sup. Fig.8

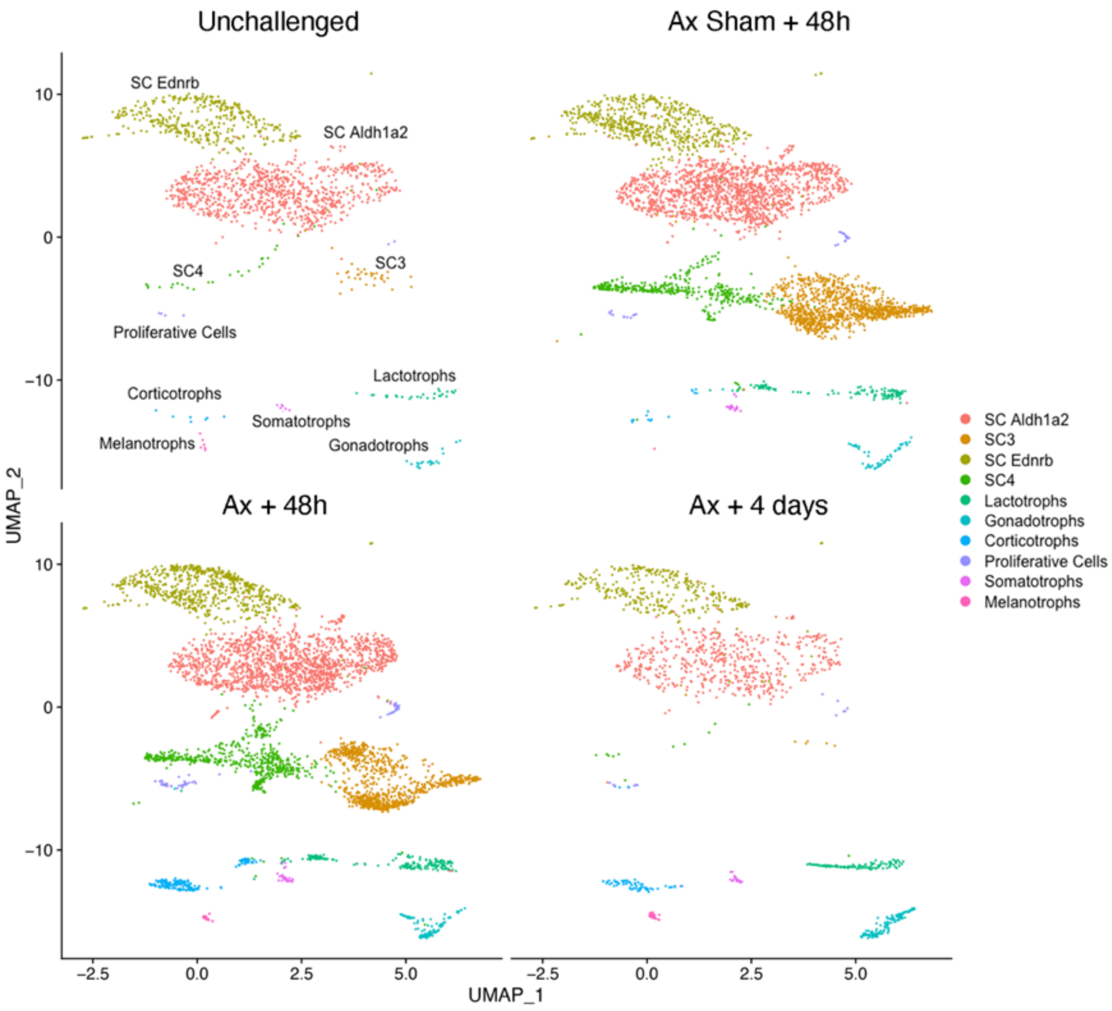

Fig. S8. UMAP clustering for filtered integrated analysis split by dataset.

A) UMAP clustering for integrated datasets from sorted SOX9iresGFP +ve cells from unchallenged, adrenalectomized and sham + 48 h, adrenalectomized + 4 days split by dataset. SC clusters 3 and 4 mostly originate from datasets collected 48 hours after surgery.

Sup. Fig.9

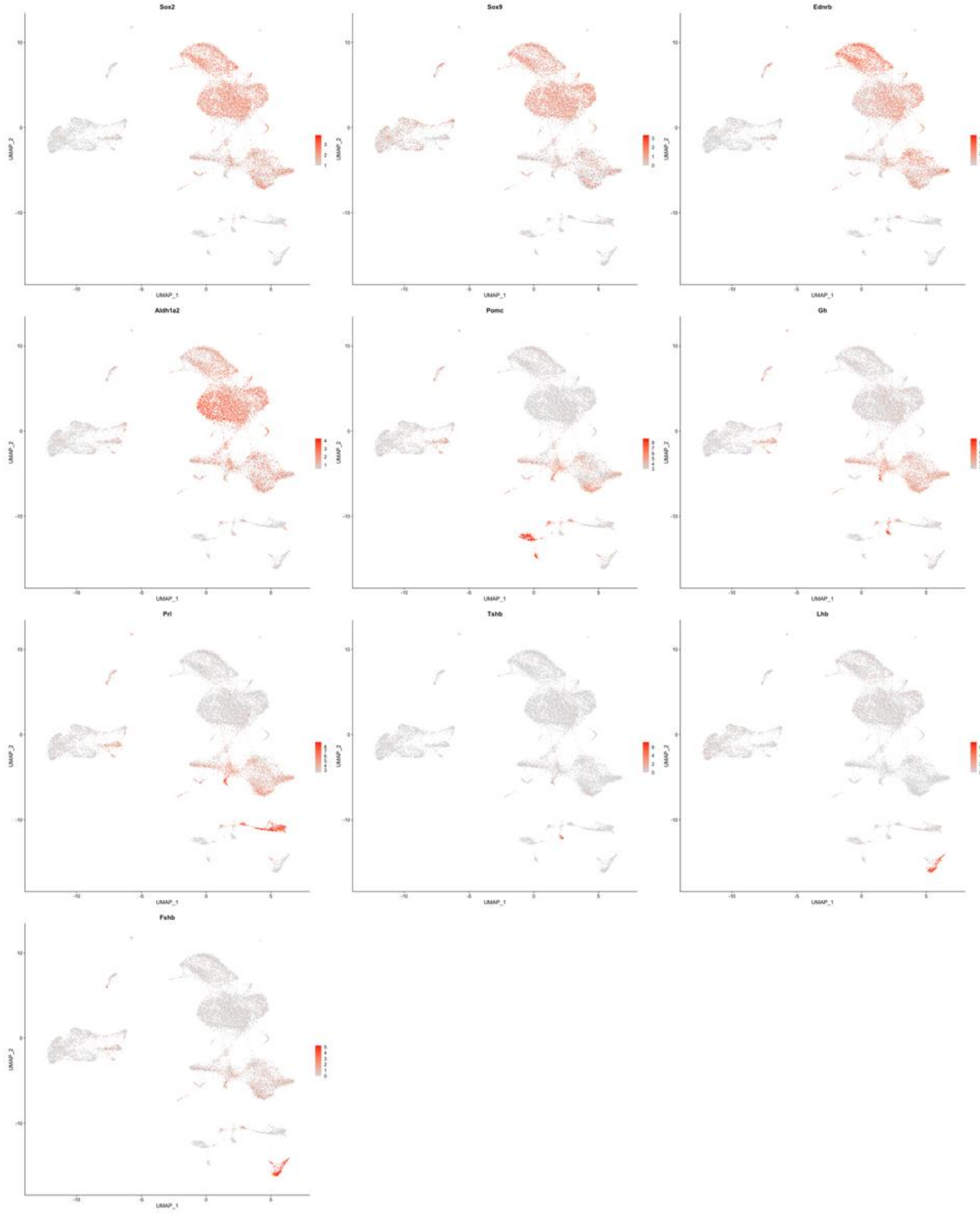
Sox2
Sox9
Ednrb
Aldh1a2
Pomc
Gh
Prl
Tshb
Lhb
Fshb
UMAP_1
UMAP_2

Fig. S9. Analyses of marker expression on UMAP clustering in Ax integrated datasets.

Markers of SC and hormone-secreting cell types are shown on the UMAP clustering for integrated datasets from GFP +ve cells sorted from *Sox9*$^{iresGFP/+}$ pituitaries from mice that were unchallenged, adrenalectomized and sham + 48 h, adrenalectomized + 4 days.

Sup. Fig.10

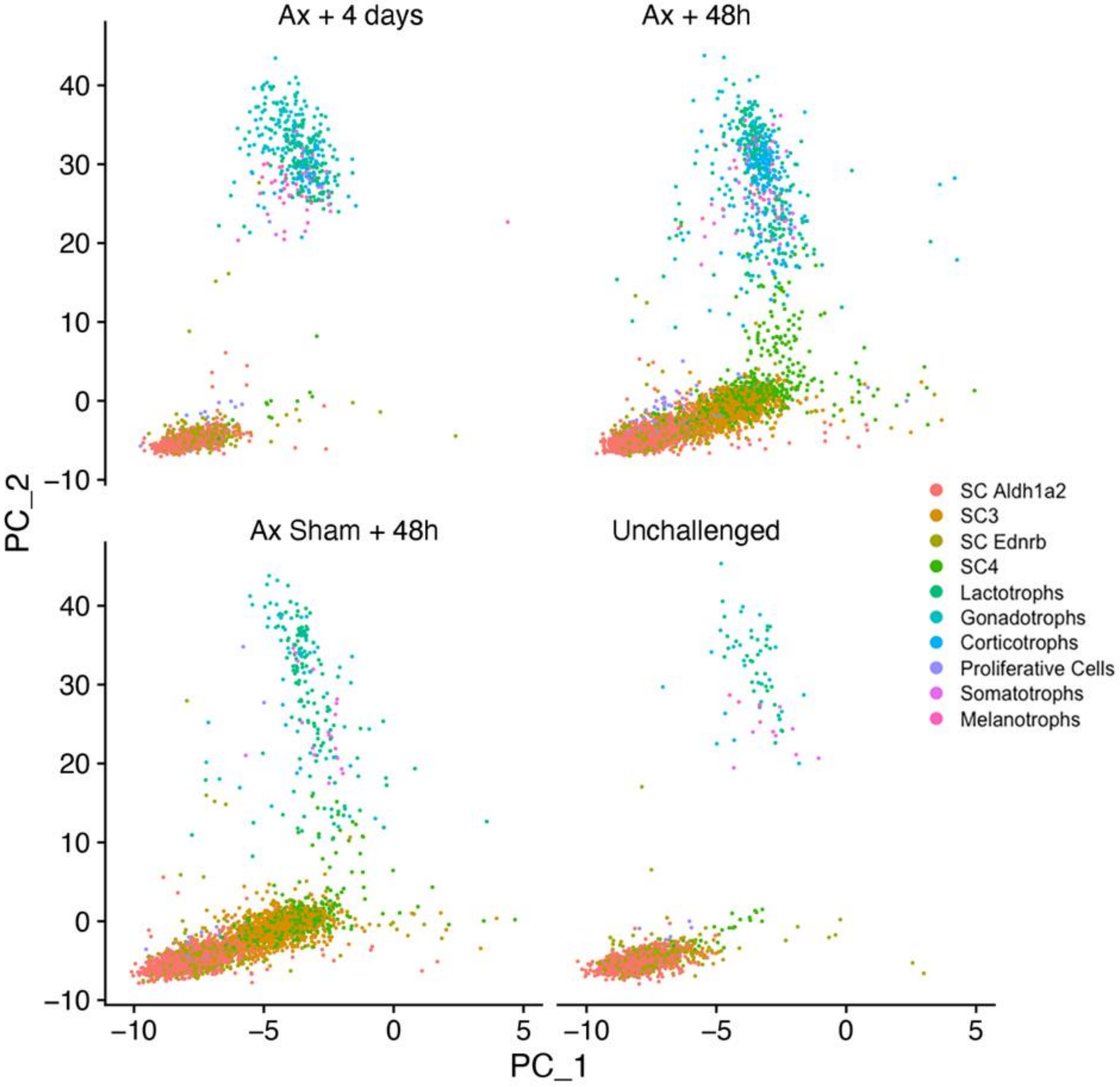

Fig. S10. PCA clustering for filtered integrated analysis split by dataset.

SC4 cluster cells located in an intermediate position between SC and endocrine cells originate mostly from Ax+48h, which is consistent with mobilization and increased commitment toward differentiation.

Sup. Fig.11

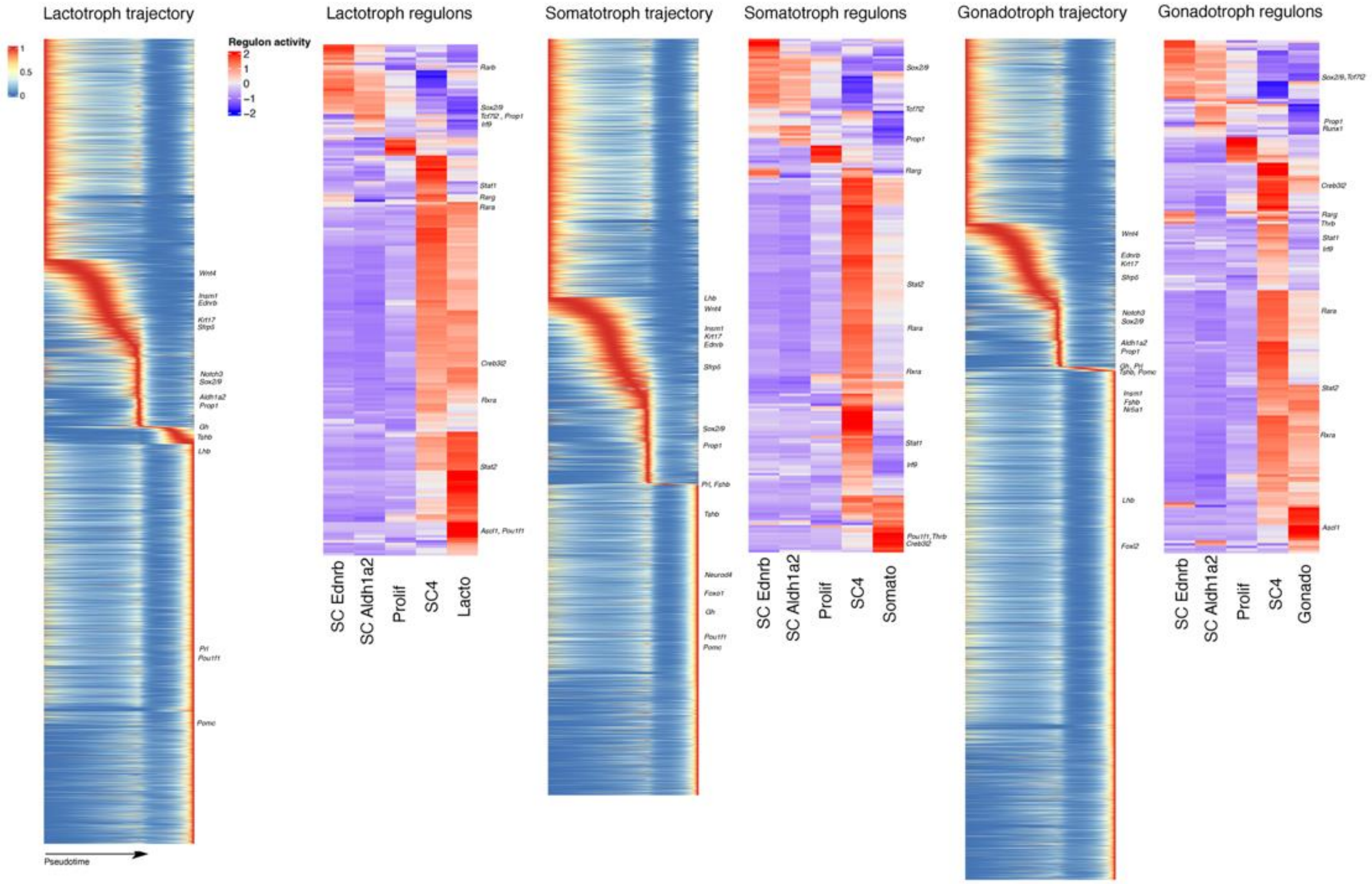

Fig. S11. Analysis of endocrine trajectories and regulons.

Heatmaps of genes associated with trajectories and SCENIC analyses for lactotroph, somatotroph and gonadotroph lineages. Representative genes are shown. Known regulators and those in common with the corticotroph analyses (Fig.5 F-H) are displayed on the heatmaps.

Sup. Fig.12

Whole pituitary dataset

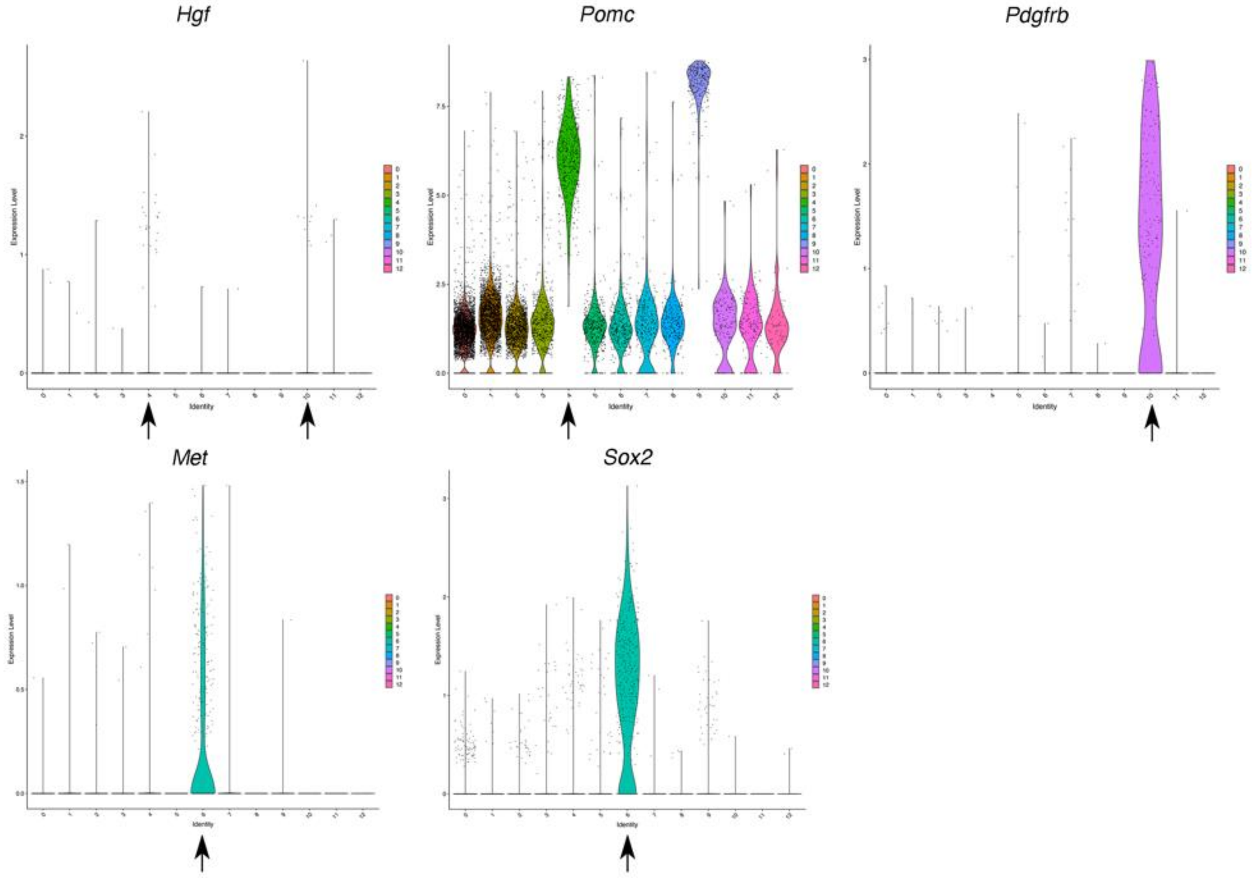

Fig. S12. Expression of *Met* and *Hgf* in the whole pituitary dataset.

Expression of *Hgf* was plotted along with *Pomc* and *Pdgfrb* suggesting expression in *Pomc* positive corticotrophs and *Pdgfrb* positive pericytes, while *Met* expression is restricted to *Sox2* positive SCs (from re-analysed dataset (*28*)). The same pattern is observed in a different dataset (*37*).

### Table S1.

**Enrichment analysis report**

Enrichment by Pathway Maps

| # | Maps | Total | pValue | Min FDR | WT Cluster 0_genelist | | | |
|---|---|---|---|---|---|---|---|---|
| | | | | | p-value | FDR | In Data | Network Objects from Active Data |
| 1 | Gamma-secretase proteolytic targets | 80 | 2.041E-11 | 2.445E-08 | 2.041E-11 | 2.445E-08 | 18 | NOTCH2 (2ICD), LRP1B, alphaAPPs, LRP1B (ICD), N-cadherin, NOTCH2 receptor, APP-C99, N-cadherin (CTF1), APP-C83 (CTF), N-cadherin (CTF2), Ephrin-B2, LRP1B (CTF), Ephrin-B2 (CTF2), APP-C59 (AICD), Amyloid beta 40, NOTCH2(NEXT), Amyloid beta 42, APP |
| 2 | Development_Extraembryonic differentiation of embryonic stem cells | 35 | 6.983E-07 | 4.183E-04 | 6.983E-07 | 4.183E-04 | 9 | ID2, ID4, HEY1, SOX2, ID1, MAP2, NCAM1, PAX6, CD9 |
| 3 | Neuroprotective action of lithium | 63 | 2.603E-06 | 7.638E-04 | 2.603E-06 | 7.638E-04 | 11 | FRAT1, Thioredoxin, Amyloid beta, P2X2, IPP-1, NR2A, WNT, NR2, p38 MAPK, Frizzled, c-Jun |
| 4 | Development_NOTCH signaling in the nervous system | 89 | 2.634E-06 | 7.638E-04 | 2.634E-06 | 7.638E-04 | 13 | NOTCH2 (2ICD), ID4, HEY1, SOX2, NOTCH2 receptor, Aquaporin 4, HES1, Adropin, Cyclin D1, NFIA, PAX6, NOTCH2(NEXT), Clusterin |
| 5 | Development_NOTCH signaling in organogenesis and embryogenesis | 77 | 3.188E-06 | 7.638E-04 | 3.188E-06 | 7.638E-04 | 12 | SIX2, NOTCH2 (2ICD), IRF6, HEY1, NOTCH2 receptor, HES1, Islet-1, HEY2, Ephrin-B2, Activin beta A, Connexin 43, NRARP |
| 6 | Inhibition of oligodendrocyte precursor cells differentiation by Wnt signaling in multiple sclerosis | 25 | 6.731E-06 | 1.220E-03 | 6.731E-06 | 1.220E-03 | 7 | FZD1, ID2, WNT9A, FZD7, WNT, Frizzled, FZD4 |
| 7 | Gamma-Secretase regulation of neuronal cell development and function | 57 | 7.131E-06 | 1.220E-03 | 7.131E-06 | 1.220E-03 | 10 | N-cadherin, Amyloid beta, APP-C99, Fe65, N-cadherin (CTF1), N-cadherin (CTF2), APP-C59 (AICD), Amyloid beta 40, betaAPPs, APP |
| 8 | Gamma-secretase regulation of gastrointestinal epithelial cell development | 28 | 1.532E-05 | 2.295E-03 | 1.532E-05 | 2.295E-03 | 7 | NOTCH2 (2ICD), HEY1, NOTCH2 receptor, HES1, HEY2, PAX6, NOTCH2(NEXT) |
| 9 | Role of IGH translocations in multiple myeloma | 40 | 2.198E-05 | 2.925E-03 | 2.198E-05 | 2.925E-03 | 8 | MafB, N-cadherin, NOTCH2 receptor, ID1, HES1, Cyclin D1, NCAM1, Histone H3 |
| 10 | Inhibition of Ephrin receptors in colorectal cancer | 30 | 2.500E-05 | 2.995E-03 | 2.500E-05 | 2.995E-03 | 7 | Ephrin-B2, Ephrin-B, WNT, Ephrin-B receptors, Ephrin-A receptors, Frizzled, Ephrin-B receptor 1 |
| 11 | Deregulation of canonical WNT signaling in major depressive disorder | 31 | 3.146E-05 | 3.193E-03 | 3.146E-05 | 3.193E-03 | 7 | FRAT1, FZD3, WNT, Cyclin D1, FZD9, MAP-1B, Frizzled |
| 12 | Transcription targets of Androgen receptor involved in Prostate Cancer | 42 | 3.198E-05 | 3.193E-03 | 3.198E-05 | 3.193E-03 | 8 | Androgen receptor, N-cadherin, ER81, Cyclin D1, Prosaposin, VIL2 (ezrin), Clusterin, APP |
| 13 | Cell adhesion_Gap junctions | 22 | 3.713E-05 | 3.357E-03 | 3.713E-05 | 3.357E-03 | 6 | Tubulin beta, Actin cytoskeletal, Actin, Calmodulin, Connexin 43, VIL2 (ezrin) |
| 14 | Stem cells_Notch signaling in medulloblastoma stem cells | 32 | 3.923E-05 | 3.357E-03 | 3.923E-05 | 3.357E-03 | 7 | NOTCH2 (2ICD), HEY1, NOTCH2 receptor, HES1, HEY2, Cyclin D1, NOTCH2(NEXT) |
| 15 | Role of activation of WNT signaling in the progression of lung cancer | 77 | 1.056E-04 | 8.437E-03 | 1.056E-04 | 8.437E-03 | 10 | FZD1, FZD3, WNT9A, FZD7, WNT, Cyclin D1, p38 MAPK, FZD9, CD147, Frizzled |
| 16 | Signal transduction_NOTCH signaling in tissue homeostasis | 81 | 1.624E-04 | 1.195E-02 | 1.624E-04 | 1.195E-02 | 10 | NOTCH2 (2ICD), Androgen receptor, HEY1, NOTCH2 receptor, HES1, HEY2, Ephrin-B2, c-Jun, OAT, CYP19 |
| 17 | WNT signaling in HCC | 40 | 1.763E-04 | 1.195E-02 | 1.763E-04 | 1.195E-02 | 7 | FZD3, Ep-CAM, FZD7, WNT, Cyclin D1, GLNA, Frizzled |
| 18 | Immune response_Induction of the antigen presentation machinery by IFN-gamma | 53 | 1.795E-04 | 1.195E-02 | 1.795E-04 | 1.195E-02 | 8 | H2-Aa, H2-Eb1, HLA-E, MHC class II, HLA-F, HLAC, CD74, HLA-DMA |
| 19 | Immune response_Antigen presentation by MHC class II | 118 | 2.422E-04 | 1.527E-02 | 2.422E-04 | 1.527E-02 | 12 | MHC class II alpha chain, HSP90, Kinesin light chain, MHC class II beta chain, Dynein 1, cytoplasmic, intermediate chains, HLA-DM, MHC class II, p38 MAPK, HSP90 beta, CLEC10A, Legumain, CD74 |
| 20 | Glutathione metabolism | 71 | 2.809E-04 | 1.683E-02 | 2.809E-04 | 1.683E-02 | 9 | GCL cat, GSTM1, GSTA3, GSTM2, GSTM5, MGST, GGTL3, GSTM1 (rodent), GSTT1 |
| 21 | Oxidative stress_Role of Sirtuin1 and PGC1-alpha in activation of antioxidant defense system | 60 | 4.308E-04 | 2.458E-02 | 4.308E-04 | 2.458E-02 | 8 | GCL cat, SCPX, Thioredoxin, PRDX1, TXNRD1, GSTM5, UCP2, PRDX5 |
| 22 | Development_TGF-beta-dependent induction of EMT via SMADs | 35 | 5.787E-04 | 3.126E-02 | 5.787E-04 | 3.126E-02 | 6 | ID2, HEY1, N-cadherin, TGF-beta 2, TGF-beta, MKL1 |
| 23 | Transcription_HIF-1 targets | 95 | 6.001E-04 | 3.126E-02 | 6.001E-04 | 3.126E-02 | 10 | ID2, SOX2, TGF-beta 2, P4HA1, ALDOC, DEC2, ENO1, REDD1, PKM2, GPI |
| 24 | Beta-catenin-dependent transcription regulation in colorectal cancer | 36 | 6.769E-04 | 3.379E-02 | 6.769E-04 | 3.379E-02 | 6 | ID2, Cyclin D1, NRCAM, LAMC2, LAMC2 (80kDa), LAMC2 (100kDa) |
| 25 | Development_NOTCH signaling activation | 82 | 8.250E-04 | 3.953E-02 | 8.250E-04 | 3.953E-02 | 9 | NOTCH2 (2ICD), HEY1, NOTCH2 receptor, HES1, HEY2, BAF60c, HDAC3, MSI1, NOTCH2(NEXT) |
| 26 | Canonical Notch signaling pathway in colorectal cancer | 52 | 9.256E-04 | 4.123E-02 | 9.256E-04 | 4.123E-02 | 7 | NOTCH2 (2ICD), NOTCH2 receptor, HES1, WNT, Cyclin D1, NOTCH2(NEXT), HES6 |
| 27 | Cell adhesion_Endothelial cell contacts by junctional mechanisms | 26 | 9.751E-04 | 4.123E-02 | 9.751E-04 | 4.123E-02 | 5 | N-cadherin, Actin cytoskeletal, Cingulin, Alpha-catenin, Connexin 43 |
| 28 | Cell adhesion_Classical cadherin-mediated cell adhesion | 26 | 9.751E-04 | 4.123E-02 | 9.751E-04 | 4.123E-02 | 5 | N-cadherin, Actin cytoskeletal, F-Actin cytoskeleton, Alpha-catenin, F-Actin |
| 29 | Cell cycle progression in Prostate Cancer | 39 | 1.050E-03 | 4.123E-02 | 1.050E-03 | 4.123E-02 | 6 | Androgen receptor, WNT, Cyclin D1, p38 MAPK, Frizzled, c-Jun |
| 30 | Activation of Notch signaling in breast cancer | 39 | 1.050E-03 | 4.123E-02 | 1.050E-03 | 4.123E-02 | 6 | HEY1, HES1, p38 MAPK, c-Jun, MSI1, Presenilin |
| 31 | Transcription_Negative regulation of HIF1A function | 69 | 1.111E-03 | 4.123E-02 | 1.111E-03 | 4.123E-02 | 8 | HSP90, PRDX2, SKP1, DEC2, HSP90 beta, KLF2, HIF-prolyl hydroxylase, EGLN2 |
| 32 | NRF2 regulation of oxidative stress response | 54 | 1.163E-03 | 4.123E-02 | 1.163E-03 | 4.123E-02 | 7 | GCL cat, GSTA3, Thioredoxin, PRDX1, TXNRD1, Actin cytoskeletal, CRM1 |
| 33 | Stem cells_NOTCH1-induced self-renewal of glioblastoma stem cells | 40 | 1.205E-03 | 4.123E-02 | 1.205E-03 | 4.123E-02 | 6 | HEY1, SOX2, ID1, HES1, HEY2, Cyclin D1 |
| 34 | Gamma-secretase regulation of osteogenesis | 40 | 1.205E-03 | 4.123E-02 | 1.205E-03 | 4.123E-02 | 6 | NOTCH2 (2ICD), HEY1, NOTCH2 receptor, HES1, Cyclin D1, NOTCH2(NEXT) |
| 35 | Stem cells_Response to hypoxia in glioblastoma stem cells | 40 | 1.205E-03 | 4.123E-02 | 1.205E-03 | 4.123E-02 | 6 | HEY1, ID1, HES1, HEY2, Cyclin D1, HIF-prolyl hydroxylase |
| 36 | DNA damage_ATM activation by DNA damage | 71 | 1.341E-03 | 4.343E-02 | 1.341E-03 | 4.343E-02 | 8 | HSP90, HMG14, p18, PP2A regulatory, HSP90 beta, Histone H2B, IHPK2, Histone H3 |
| 37 | Development_Positive regulation of STK3/4 (Hippo) pathway and negative regulation of YAP/TAZ func | 71 | 1.341E-03 | 4.343E-02 | 1.341E-03 | 4.343E-02 | 8 | EBP50, Angiomotin (AMOT), Actin cytoskeletal, MALS-3, Alpha-catenin, Alpha-1 catenin, G-protein alpha-s, 14-3-3 |
| 38 | G-protein signaling_Rac3 regulation pathway | 17 | 1.453E-03 | 4.580E-02 | 1.453E-03 | 4.580E-02 | 4 | Androgen receptor, Actin cytoplasmic 2, Calmyrin, F-Actin |
| 39 | Ubiquinone metabolism | 74 | 1.756E-03 | 5.333E-02 | 1.756E-03 | 5.333E-02 | 8 | MTND1, NDUFA4, MT-ND5, MTND4, MTND2, NDUFC2, DAP13, MTND3 |
| 40 | Notch signaling in breast cancer | 58 | 1.781E-03 | 5.333E-02 | 1.781E-03 | 5.333E-02 | 7 | NOTCH2 (2ICD), HEY1, NOTCH2 receptor, HES1, HEY2, Cyclin D1, NOTCH2(NEXT) |
| 41 | Androgen receptor activation and downstream signaling in Prostate cancer | 110 | 1.865E-03 | 5.450E-02 | 1.865E-03 | 5.450E-02 | 10 | Androgen receptor, N-cadherin, ER81, SGK1, Cyclin D1, Prosaposin, FGF1, VIL2 (ezrin), Clusterin, APP |
| 42 | Development_Transcription factors in segregation of hepatocytic lineage | 30 | 1.912E-03 | 5.454E-02 | 1.912E-03 | 5.454E-02 | 5 | Activin A, Activin B, Cyclin D1, Activin, GGTL3 |
| 43 | Aminoglycoside- and cisplatin-induced hair cell death | 112 | 2.134E-03 | 5.749E-02 | 2.134E-03 | 5.749E-02 | 10 | NOL3, AP-1, p38 MAPK, HDAC3, Histone H2B, Cathepsin D, CX3CL1, c-Jun, Histone H3, LMO4 |
| 44 | Role of inhibition of WNT signaling in the progression of lung cancer | 31 | 2.224E-03 | 5.749E-02 | 2.224E-03 | 5.749E-02 | 5 | N-cadherin, CD82, FZD9, Frizzled, c-Jun |
| 45 | Development_Embryonal epaxial myogenesis | 31 | 2.224E-03 | 5.749E-02 | 2.224E-03 | 5.749E-02 | 5 | FZD1, HES1, WNT, Pitx2, Frizzled |
| 46 | Development_Negative regulation of WNT/Beta-catenin signaling at the receptor level | 45 | 2.251E-03 | 5.749E-02 | 2.251E-03 | 5.749E-02 | 6 | LRP4, Amyloid beta, WNT, Frizzled, FZD4, Syndecans |
| 47 | Inhibition of WNT5A-dependent non-canonical pathway in colorectal cancer | 19 | 2.255E-03 | 5.749E-02 | 2.255E-03 | 5.749E-02 | 4 | Cyclin D1, Calmodulin, CRM1, c-Jun |
| 48 | Neurophysiological process_ACM1, ACM3 and ACM5 signaling in the brain | 78 | 2.461E-03 | 6.141E-02 | 2.461E-03 | 6.141E-02 | 8 | alphaAPPs, TRPM3, APP-C99, APP-C83 (CTF), Calmodulin, betaAPPs, CACNB3, APP |
| 49 | Inhibition of remyelination in multiple sclerosis: role of cell-cell and ECM-cell interactions | 47 | 2.820E-03 | 6.894E-02 | 2.820E-03 | 6.894E-02 | 6 | CD81, Actin cytoskeletal, CD82, NCAM1, Connexin 43, CD9 |
| 50 | Role of SHH and Notch in SCLC | 33 | 2.956E-03 | 6.944E-02 | 2.956E-03 | 6.944E-02 | 5 | NOTCH2 (2ICD), NOTCH2 receptor, HES1, AL1A1, NOTCH2(NEXT) |

**Enrichment analysis report**

Enrichment by Pathway Maps

| # | Maps | Total | pValue | Min FDR | WT Cluster 2_genelist p-value | FDR | In Data | Network Objects from Active Data |
|---|---|---|---|---|---|---|---|---|
| 1 | Cytoskeleton remodeling_Keratin filaments | 36 | 5.894E-09 | 4.697E-06 | 5.894E-09 | 4.697E-06 | 8 | Keratin 19, Keratin 17, Tubulin alpha, Keratin 8, Keratin 8/18, Vimentin, Keratin 18, Keratin 7 |
| 2 | Cytoskeleton remodeling_Regulation of actin cytoskeleton organization by the kinase effectors of Rho | 58 | 3.015E-07 | 1.202E-04 | 3.015E-07 | 1.202E-04 | 8 | Rac3, MLCP (cat), Rhov, Spectrin, MRLC, Rac1-related, WRCH-1, Cdc42 subfamily |
| 3 | Cell adhesion_Desmosomes | 19 | 1.935E-06 | 5.139E-04 | 1.935E-06 | 5.139E-04 | 5 | Keratin 17, Keratin 8/18, Vimentin, Keratin 18, Plakoglobin |
| 4 | Cell adhesion_Tight junctions | 44 | 1.038E-05 | 1.891E-03 | 1.038E-05 | 1.891E-03 | 6 | PKC-zeta, Tubulin alpha, Occludin, MRLC, CRB3, ZO-3 |
| 5 | Mechanisms of resistance to EGFR inhibitors in lung cancer | 45 | 1.186E-05 | 1.891E-03 | 1.186E-05 | 1.891E-03 | 6 | HSP90, Claudin-7, Ep-CAM, Vimentin, Claudin-4, TACSTD2 (TROP2) |
| 6 | Canonical Notch signaling pathway in colorectal cancer | 52 | 2.775E-05 | 3.687E-03 | 2.775E-05 | 3.687E-03 | 6 | KLF4, NOTCH1 (NICD), NOTCH1 (NEXT), KLF5, NOTCH1 receptor, NOTCH1 precursor |
| 7 | Oxidative stress_ROS-induced cellular signaling | 108 | 3.412E-05 | 3.885E-03 | 3.412E-05 | 3.885E-03 | 8 | HSP27, NOTCH1 (NICD), TXNIP (VDUP1), FTH1, NFKBIA, Pin1, DLC1 (Dynein LC8a), PKC |
| 8 | Possible regulation of HSF-1/ chaperone pathway in Huntington's disease | 21 | 8.687E-05 | 8.655E-03 | 8.687E-05 | 8.655E-03 | 4 | HSP90, HSP27, PLA2, HSP90 alpha |
| 9 | Signal transduction_Relaxin family peptides signaling via RXFP1 and RXFP2 receptors | 70 | 1.508E-04 | 1.335E-02 | 1.508E-04 | 1.335E-02 | 6 | NOTCH1 (NICD), PKC-zeta, NOTCH1 (NEXT), EDNRB, NFKBIA, I-kB |
| 10 | Cytoskeleton remodeling_Regulation of actin cytoskeleton nucleation and polymerization by Rho GTP | 46 | 1.776E-04 | 1.415E-02 | 1.776E-04 | 1.415E-02 | 5 | Rac3, mDIA2(DIAPH3), Rac1-related, DRF, Cdc42 subfamily |
| 11 | Cell adhesion_Endothelial cell contacts by junctional mechanisms | 26 | 2.075E-04 | 1.503E-02 | 2.075E-04 | 1.503E-02 | 4 | Occludin, Vimentin, Claudin-3, Plakoglobin |
| 12 | Glucocorticoids-mediated inhibition of pro-constrictory and pro-inflammatory signaling in airway smo | 49 | 2.403E-04 | 1.596E-02 | 2.403E-04 | 1.596E-02 | 5 | MLCP (cat), PLA2, NFKBIA, MRLC, MKP-1 |
| 13 | Notch signaling in oligodendrocyte precursor cell differentiation in multiple sclerosis | 29 | 3.209E-04 | 1.968E-02 | 3.209E-04 | 1.968E-02 | 4 | NOTCH1 (NICD), NOTCH1 (NEXT), SOX10, NOTCH1 receptor |
| 14 | Development_Auditory hair cell differentiation in embryogenesis | 31 | 4.176E-04 | 2.378E-02 | 4.176E-04 | 2.378E-02 | 4 | NOTCH1 (NICD), NOTCH1 (NEXT), NOTCH1 receptor, NOTCH1 precursor |
| 15 | Skeletal muscle atrophy in COPD | 57 | 4.901E-04 | 2.604E-02 | 4.901E-04 | 2.604E-02 | 5 | NFKBIA, I-kB, Vimentin, 14-3-3, JunD |
| 16 | Notch signaling in breast cancer | 58 | 5.314E-04 | 2.620E-02 | 5.314E-04 | 2.620E-02 | 5 | NOTCH1 (NICD), NOTCH1 (NEXT), Pin1, NOTCH1 receptor, NOTCH1 precursor |
| 17 | Development_NOTCH signaling in the nervous system | 89 | 5.589E-04 | 2.620E-02 | 5.589E-04 | 2.620E-02 | 6 | NOTCH1 (NICD), PKC-zeta, NOTCH1 (NEXT), SOX10, NOTCH1 receptor, Clusterin |
| 18 | Development_WNT and Notch signaling in early cardiac myogenesis | 35 | 6.706E-04 | 2.969E-02 | 6.706E-04 | 2.969E-02 | 4 | NOTCH1 (NICD), SFRP5, NOTCH1 (NEXT), NOTCH1 receptor |
| 19 | Immune response_IL-4 signaling pathway | 94 | 7.471E-04 | 2.995E-02 | 7.471E-04 | 2.995E-02 | 6 | AP-1, PKC-zeta, Tubulin alpha, NFKBIA, PKC, JunD |
| 20 | Cytoskeleton remodeling_Substance P mediated membrane blebbing | 16 | 7.515E-04 | 2.995E-02 | 7.515E-04 | 2.995E-02 | 3 | MLCP (cat), Tubulin alpha, MRLC |
| 21 | Stem cells_CD30 signaling in transformed embryonic stem cells | 18 | 1.077E-03 | 4.087E-02 | 1.077E-03 | 4.087E-02 | 3 | KLF4, I-kB, TRAF1 |
| 22 | Development_NOTCH-induced EMT | 19 | 1.268E-03 | 4.593E-02 | 1.268E-03 | 4.593E-02 | 3 | NOTCH1 (NICD), NOTCH1 (NEXT), NOTCH1 receptor |
| 23 | Signal transduction_Non-neuronal ACM1, ACM3 and ACM5 signaling | 74 | 1.613E-03 | 5.588E-02 | 1.613E-03 | 5.588E-02 | 5 | MLCP (cat), PKC-zeta, MRLC2, MRLC, PKC |
| 24 | Role of DNA methylation in progression of multiple myeloma | 45 | 1.746E-03 | 5.797E-02 | 1.746E-03 | 5.797E-02 | 4 | AP-1, SFRP5, DKK3, NOTCH1 receptor |
| 25 | Signal transduction_Angiotensin II/AGTR1 signaling via Notch, Beta-catenin and NF-kB pathways | 78 | 2.037E-03 | 6.495E-02 | 2.037E-03 | 6.495E-02 | 5 | NOTCH1 (NICD), NOTCH1 (NEXT), I-kB, NOTCH1 receptor, PKC |
| 26 | Development_NOTCH signaling activation | 82 | 2.538E-03 | 7.781E-02 | 2.538E-03 | 7.781E-02 | 5 | NOTCH1 (NICD), PKC-zeta, NOTCH1 (NEXT), NOTCH1 receptor, NOTCH1 precursor |
| 27 | HSP70 and HSP40-dependent folding in Huntington's disease | 25 | 2.861E-03 | 8.446E-02 | 2.861E-03 | 8.446E-02 | 3 | HSP90, HSP27, HSP90 alpha |
| 28 | Signal transduction_Additional pathways of NF-kB activation (in the cytoplasm) | 52 | 2.981E-03 | 8.485E-02 | 2.981E-03 | 8.485E-02 | 4 | PKC-zeta, NFKBIA, Pin1, I-kB |
| 29 | Gamma-secretase regulation of gastrointestinal epithelial cell development | 28 | 3.974E-03 | 1.056E-01 | 3.974E-03 | 1.056E-01 | 3 | NOTCH1 (NICD), NOTCH1 (NEXT), NOTCH1 receptor |
| 30 | Rho-dependent regulation of normal and asthmatic smooth muscle contraction | 28 | 3.974E-03 | 1.056E-01 | 3.974E-03 | 1.056E-01 | 3 | MLCP (cat), MRLC, PKC |
| 31 | Development_VEGF signaling via VEGFR2 - generic cascades | 93 | 4.372E-03 | 1.070E-01 | 4.372E-03 | 1.070E-01 | 5 | HSP90, HSP27, I-kB, PKC, PLA2G5 |
| 32 | Cigarette smoke-mediated regulation of NRF2-antioxidant pathway in airway epithelial cells | 29 | 4.396E-03 | 1.070E-01 | 4.396E-03 | 1.070E-01 | 3 | GSTO1, TALDO, DJ-1 |
| 33 | Aberrant production of IL-2 and IL-17 in SLE T cells | 58 | 4.430E-03 | 1.070E-01 | 4.430E-03 | 1.070E-01 | 4 | NOTCH1 (NICD), AP-1, I-kB, NOTCH1 precursor |
| 34 | Gamma-secretase regulation of angiogenesis | 30 | 4.843E-03 | 1.103E-01 | 4.843E-03 | 1.103E-01 | 3 | NOTCH1 (NICD), NOTCH1 (NEXT), NOTCH1 receptor |
| 35 | Signal transduction_Additional pathways of NF-kB activation (in the nucleus) | 30 | 4.843E-03 | 1.103E-01 | 4.843E-03 | 1.103E-01 | 3 | PKC-zeta, NFKBIA, I-kB |
| 36 | Role of inhibition of WNT signaling in the progression of lung cancer | 31 | 5.317E-03 | 1.177E-01 | 5.317E-03 | 1.177E-01 | 3 | Keratin 8, Vimentin, Keratin 18 |
| 37 | Stem cells_Notch signaling in medulloblastoma stem cells | 32 | 5.818E-03 | 1.253E-01 | 5.818E-03 | 1.253E-01 | 3 | NOTCH1 (NICD), NOTCH1 (NEXT), NOTCH1 receptor |
| 38 | Role of SHH and Notch in SCLC | 33 | 6.347E-03 | 1.292E-01 | 6.347E-03 | 1.292E-01 | 3 | NOTCH1 (NICD), NOTCH1 (NEXT), NOTCH1 receptor |
| 39 | LRRK2 in neurons in Parkinson's disease | 33 | 6.347E-03 | 1.292E-01 | 6.347E-03 | 1.292E-01 | 3 | HSP90, PKC-zeta, 14-3-3 |
| 40 | Development_NOTCH1 signaling in the epidermis | 65 | 6.645E-03 | 1.292E-01 | 6.645E-03 | 1.292E-01 | 4 | NOTCH1 (NICD), NOTCH1 (NEXT), I-kB, NOTCH1 receptor |
| 41 | Signal transduction_PDGF signaling via MAPK cascades | 65 | 6.645E-03 | 1.292E-01 | 6.645E-03 | 1.292E-01 | 4 | HSP27, KLF4, AP-1, JunD |
| 42 | Development_Epigenetic and transcriptional regulation of oligodendrocyte precursor cell differentiation | 34 | 6.903E-03 | 1.310E-01 | 6.903E-03 | 1.310E-01 | 3 | NOTCH1 (NICD), SOX10, NOTCH1 receptor |
| 43 | Development_TGF-beta-dependent induction of EMT via SMADs | 35 | 7.488E-03 | 1.356E-01 | 7.488E-03 | 1.356E-01 | 3 | NOTCH1 (NICD), Occludin, Vimentin |
| 44 | Immune response_Lipoxins and Resolvin E1 inhibitory action on neutrophil functions | 35 | 7.488E-03 | 1.356E-01 | 7.488E-03 | 1.356E-01 | 3 | PKC-zeta, NFKBIA, I-kB |
| 45 | HBV signaling via protein kinases leading to HCC | 36 | 8.100E-03 | 1.435E-01 | 8.100E-03 | 1.435E-01 | 3 | AP-1, Pin1, PKC |
| 46 | Immune response_Plasmin signaling | 70 | 8.612E-03 | 1.492E-01 | 8.612E-03 | 1.492E-01 | 4 | Annexin II, NFKBIA, PKC, JunD |
| 47 | TNF-alpha-induced inflammatory signaling in normal and asthmatic airway epithelium | 38 | 9.412E-03 | 1.576E-01 | 9.412E-03 | 1.576E-01 | 3 | NFKBIA, FN14(TNFRSF12A), I-kB |
| 48 | Signal transduction_Calcium-mediated signaling | 72 | 9.494E-03 | 1.576E-01 | 9.494E-03 | 1.576E-01 | 4 | MLCP (cat), I-kB, PKC, 14-3-3 |
| 49 | Activation of Notch signaling in breast cancer | 39 | 1.011E-02 | 1.630E-01 | 1.011E-02 | 1.630E-01 | 3 | KLF4, NOTCH1 (NICD), NOTCH1 precursor |
| 50 | Gamma-secretase regulation of osteogenesis | 40 | 1.084E-02 | 1.630E-01 | 1.084E-02 | 1.630E-01 | 3 | NOTCH1 (NICD), NOTCH1 (NEXT), NOTCH1 receptor |

Pathway analysis for clusters 0 and 2.

Gene markers for WT Cluster 0 were used to determine enriched pathways with Metacore using a hypergeometric test. Pathways with an adjusted p value < 0.05 are shown in the table. Genes present in our dataset are highlighted in the top5 pathways.

#### Table S2.

| | | Number of cells | | | |
|---|---|---|---|---|---|
| ID | Cluster ID | AX+ 4days | AX+ 48h00 | AXSham | WT |
| SC Aldh1a2 | 0 | 541 | 1606 | 1492 | 910 |
| SC3 | 1 | 7 | 941 | 1068 | 36 |
| SC Ednrb | 2 | 334 | 685 | 593 | 409 |
| Dcn | 3 | 427 | 403 | 493 | 484 |
| SC4 | 4 | 12 | 603 | 387 | 27 |
| Dcn/vasc | 5 | 7 | 212 | 237 | 24 |
| Lacto | 6 | 127 | 198 | 120 | 28 |
| Gonado | 7 | 136 | 94 | 56 | 22 |
| Cortico | 8 | 69 | 199 | 14 | 7 |
| Dcn/vasc | 9 | 22 | 72 | 86 | 21 |
| Prolif | 10 | 13 | 69 | 26 | 6 |
| Somato | 11 | 24 | 37 | 18 | 7 |
| Melano | 12 | 31 | 33 | 1 | 6 |
| Dcn/vasc | 13 | 0 | 25 | 13 | 5 |
| Dcn/vasc | 14 | 15 | 18 | 4 | 1 |
| | | 1765 | 5195 | 4608 | 1993 |
| Endo cells total | | 387 | 561 | 209 | 70 |
| %endo/total (endo=Clusters 6,7,8,14 and15) | | 22 | 11 | 4.5 | 3.5 |
| % lacto/endo | | 32 | 35 | 57 | 40 |
| % cortico/endo | | 18 | 35 | 7 | 3 |
| | | AX+ 4days | AX+ 48h00 | AXSham | WT |
| %cortico/total | | 4 | 3.8 | 0.3 | 0.3 |
| %lacto/total | | 7.2 | 3.8 | 2.6 | 1.4 |
| % somato/total | | 1.3 | 0.7 | 0.4 | 0.3 |
| % gonado/total | | 7.7 | 1.8 | 1.2 | 1.1 |

<u>Endocrine cell cluster numbers from Ax integrated dataset (Fig.5)</u>

**Table S3.**

| Showing 1 to 6 of 6 entries, 5 total columns | | | | | | |
|---|---|---|---|---|---|---|
| Pairwise for proportion test (Fig.5B) | | | | | | |
| | **group1** | **group2** | **p** | **p.adj** | **p.adj.signif** | |
| **1** | SC Aldh1a2 | SC3 | 5.35E-04 | 1.02E-02 | * | |
| **2** | SC3 | SC Ednrb | 1.82E-04 | 4.18E-03 | ** | |
| **3** | SC Aldh1a2 | SC4 | 7.50E-07 | 2.03E-05 | **** | |
| **4** | SC3 | SC4 | 5.55E-13 | 1.72E-11 | **** | |
| **5** | SC Ednrb | SC4 | 5.76E-04 | 1.04E-02 | * | |
| **6** | SC Aldh1a2 | Prolif | 9.91E-05 | 2.38E-03 | ** | |
| **7** | SC3 | Prolif | 1.50E-06 | 3.74E-05 | **** | |
| **8** | SC Ednrb | Prolif | 4.83E-04 | 1.01E-02 | * | |
| **9** | SC Aldh1a2 | Lacto | 4.88E-04 | 1.01E-02 | * | |
| **10** | SC3 | Lacto | 4.36E-07 | 1.22E-05 | **** | |
| **11** | SC3 | Gonado | 2.54E-04 | 5.59E-03 | ** | |
| **12** | SC Aldh1a2 | Cortico | 1.02E-31 | 3.56E-30 | **** | |
| **13** | SC3 | Cortico | 7.30E-38 | 2.63E-36 | **** | |
| **14** | SC Ednrb | Cortico | 1.46E-27 | 4.97E-26 | **** | |
| **15** | SC4 | Cortico | 1.39E-19 | 4.60E-18 | **** | |
| **16** | Prolif | Cortico | 1.36E-06 | 3.54E-05 | **** | |
| **17** | Lacto | Cortico | 1.23E-15 | 3.95E-14 | **** | |
| **18** | Gonado | Cortico | 6.99E-13 | 2.10E-11 | **** | |
| **19** | Cortico | Somato | 3.40E-07 | 9.87E-06 | **** | |
| Showing 1 to 19 of 19 entries, 5 total columns | | | | | | |
| Pairwise for proportion test (Fig.6B) | | | | | | |
| | n | Statistic | df | p.value | adj.p.value | p.signif |
| POMC.in.ma | 27347 | 174.098914 | 1 | 9.42E-40 | 2.51E-39 | **** |
| PRL.in.males | 35836 | 15.302442 | 1 | 9.16E-05 | 1.47E-04 | **** |
| LH.in.males | 35836 | 1.803625 | 1 | 1.79E-01 | 1.79E-01 | ns |
| GH.in.males | 31383 | 10.574053 | 1 | 1.15E-03 | 1.31E-03 | ** |
| POMC.in.fem | 60958 | 404.277391 | 1 | 6.45E-90 | 5.16E-89 | **** |
| PRL.in.femal | 62693 | 235.560711 | 1 | 3.65E-53 | 1.46E-52 | **** |
| LH.in.female | 62693 | 10.638566 | 1 | 1.11E-03 | 1.31E-03 | ** |
| GH.in.female | 57634 | 165.802853 | 1 | 6.11E-38 | 1.22E-37 | **** |

<u>Pairwise for proportion test results.</u>

**Table S4**

| SOX2; AQP3 parenchyma countings | | | | | | |
|---|---|---|---|---|---|---|
| | SOX2 | SOX2 only | | Deduced SOX2;AQP3 | | % SOX2;AQP |
| XY.1 | 668 | 228 | | 440 | | 65.8682635 |
| XY.2 | 486 | 210 | | 276 | | 56.7901235 |
| XY.3 | 522 | 128 | | 394 | | 75.4789272 |

**Fig.2 countings**

| | | | | | | | | |
|---|---|---|---|---|---|---|---|---|
| Sox9;PDGFRb counts were exclusively performed in the parenchyma (cleft excluded) on triple immunofluorescence (GFP fo | | | | | | | | |
| Manual coutings | | | | | | | | |
| Animal | Sex | Age | Sox9iresGFP;Sox9 | | Sox9iresGFP;Sox9;PDGFRb | | | % Sox9iresGFP;S |
| 1 | XY | 6m-old | 387 | | 39 | | | 10.0775194 |
| 2 | XY | 6 m-old | 265 | | 19 | | | 7.16981132 |
| 3 | XY | 6 m-old | 291 | | 24 | | | 8.24742268 |
| 4 | XY | 8 m-old | 616 | | 39 | | | 6.33116883 |
| 1 | XX | 2 m-old | 331 | | 7 | | | 2.11480363 |
| 2 | XX | 4 m-old | 734 | | 10 | | | 1.36239782 |
| 3 | XX | 8 m-old | 739 | | 9 | | | 1.21786198 |
| 4 | XX | 4 m-old | 807 | | 5 | | | 0.61957869 |

| | | | | | | | |
|---|---|---|---|---|---|---|---|
| Cell countings after lineage tracing using Wnt1Cre on triple immunofluorescence (GFP for Wnt1Cre;R26 eYFP, Sox9 and PD | | | | | | | |
| Manual countings | | | | | | | |
| Animal | Sex | Age | Sox9;PDGFRb | | Sox9;PDGFRb;eYFP | | % Sox9;PDGFRb;eYFP/Sox9;eY |
| 1 | XY | 6w-old | 49 | | 36 | | 73.4693878 |
| 2 | XY | 6w-old | 55 | | 43 | | 78.1818182 |
| 3 | XY | 6w-old | 35 | | 26 | | 74.2857143 |
| 4 | XY | 6w-old | 33 | | 25 | | 75.7575758 |
| 5 | XY | 6w-old | 55 | | 43 | | 78.1818182 |
| 6 | XY | 6w-old | 66 | | 62 | | 93.9393939 |
| 1 | XX | 6w-old | 21 | | 9 | | 42.8571429 |
| 2 | XX | 6w-old | 17 | | 9 | | 52.9411765 |
| 3 | XX | 6w-old | 19 | | 11 | | 57.8947368 |

**Fig.3 countings**

| | | | | | | |
|---|---|---|---|---|---|---|
| Dissociated Sox9iresGFP pituitaries were FACSorted and plated 4 days after surgery. | | | | | | |
| Immunofluorescent stainings: GH and POMC separately, LH and PRL as a double-immuno. | | | | | | |
| Automated countings | | | | | | |
| Experiment | positive | DAPI | sum | Hormone | Sex | % Horm/DAF |
| Ax | 20 | 2617 | 2637 | POMC | Male | 0.75843762 |
| Ax | 162 | 5878 | 6040 | POMC | Male | 2.68211921 |
| Ax | 221 | 4945 | 5166 | POMC | Male | 4.27797135 |
| ShAx | 5 | 3255 | 3260 | POMC | Male | 0.15337423 |
| ShAx | 25 | 4183 | 4208 | POMC | Male | 0.59410646 |
| ShAx | 72 | 5964 | 6036 | POMC | Male | 1.19284294 |
| Ax | 33 | 4224 | 4257 | POMC | Female | 0.7751938 |
| Ax | 276 | 10518 | 10794 | POMC | Female | 2.5569761 |
| Ax | 479 | 10744 | 11223 | POMC | Female | 4.26802103 |
| ShAx | 17 | 6684 | 6701 | POMC | Female | 0.25369348 |
| ShAx | 251 | 17558 | 17809 | POMC | Female | 1.40939974 |
| ShAx | 20 | 10154 | 10174 | POMC | Female | 0.19657952 |
| Ax | 9 | 2836 | 2845 | PRL | Male | 0.31634446 |
| Ax | 142 | 7180 | 7322 | PRL | Male | 1.93936083 |
| Ax | 165 | 7214 | 7379 | PRL | Male | 2.23607535 |
| ShAx | 8 | 3439 | 3447 | PRL | Male | 0.23208587 |
| ShAx | 119 | 4631 | 4750 | PRL | Male | 2.50526316 |
| ShAx | 312 | 9781 | 10093 | PRL | Male | 3.09125136 |
| Ax | 48 | 3331 | 3379 | PRL | Female | 1.42053862 |
| Ax | 1093 | 12121 | 13214 | PRL | Female | 8.2715302 |
| Ax | 415 | 13231 | 13646 | PRL | Female | 3.04118423 |
| ShAx | 50 | 4512 | 4562 | PRL | Female | 1.09601052 |
| ShAx | 764 | 17302 | 18066 | PRL | Female | 4.22893834 |
| ShAx | 83 | 9743 | 9826 | PRL | Female | 0.84469774 |
| Ax | 10 | 2835 | 2845 | LH | Male | 0.35149385 |
| Ax | 156 | 7166 | 7322 | LH | Male | 2.13056542 |
| Ax | 183 | 7196 | 7379 | LH | Male | 2.48001084 |
| ShAx | 10 | 3437 | 3447 | LH | Male | 0.29010734 |
| ShAx | 110 | 4640 | 4750 | LH | Male | 2.31578947 |
| ShAx | 282 | 9811 | 10093 | LH | Male | 2.79401565 |
| Ax | 12 | 3367 | 3379 | LH | Female | 0.35513466 |
| Ax | 153 | 13061 | 13214 | LH | Female | 1.15786287 |
| Ax | 87 | 13559 | 13646 | LH | Female | 0.63754947 |
| ShAx | 10 | 4552 | 4562 | LH | Female | 0.2192021 |
| ShAx | 130 | 17936 | 18066 | LH | Female | 0.71958375 |
| ShAx | 58 | 9768 | 9826 | LH | Female | 0.59027071 |
| Ax | 53 | 2721 | 2774 | GH | Male | 1.91059841 |
| Ax | 196 | 6195 | 6391 | GH | Male | 3.06681271 |
| Ax | 291 | 7313 | 7604 | GH | Male | 3.82693319 |
| ShAx | 46 | 2823 | 2869 | GH | Male | 1.60334611 |
| ShAx | 178 | 6409 | 6587 | GH | Male | 2.70229239 |
| ShAx | 155 | 5003 | 5158 | GH | Male | 3.00504071 |
| Ax | 19 | 2331 | 2350 | GH | Female | 0.80851064 |
| Ax | 923 | 11164 | 12087 | GH | Female | 7.63630347 |
| Ax | 571 | 9587 | 10158 | GH | Female | 5.62118527 |
| ShAx | 36 | 3430 | 3466 | GH | Female | 1.03866128 |
| ShAx | 1034 | 19796 | 20830 | GH | Female | 4.96399424 |
| ShAx | 194 | 8549 | 8743 | GH | Female | 2.21891799 |

**Figure 6B countings**

| | | | | |
|---|---|---|---|---|
| A week after surgeries Sox9iresCreERT2;R26eYFP pituitaries were processed for immunofluorescence on sections | | | | |
| Countings were performed manually on sections | | | | |
| Animal | Age | Sex | Surgery | ACTH;eYFP |
| 1 | <=7 w-old | XX | Ax | 66 |
| 2 | <=7 w-old | XX | Ax | 34 |
| 3 | <=7 w-old | XX | Ax | 171 |
| 4 | <=7 w-old | XX | Ax | 19 |
| 1 | <=7 w-old | XY | Ax | 84 |
| 2 | <=7 w-old | XY | Ax | 93 |
| 3 | <=7 w-old | XY | Ax | 66 |
| 4 | <=7 w-old | XY | Ax | 47 |
| 5 | <=7 w-old | XY | Ax | 50 |
| 6 | <=7 w-old | XY | Ax | 78 |
| Animal | Age | Sex | Surgery | PRL;eYFP |
| 1 | <=7 w-old | XX | Ax | 38 |
| 2 | <=7 w-old | XX | Ax | 68 |
| 3 | <=7 w-old | XX | Ax | 44 |
| 4 | <=7 w-old | XX | Ax | 69 |
| 5 | <=7 w-old | XX | Ax | 86 |
| 1 | <=7 w-old | XY | Ax | 1 |
| 2 | <=7 w-old | XY | Ax | 1 |
| 3 | <=7 w-old | XY | Ax | 0 |
| 4 | <=7 w-old | XY | XY | 0 |
| Animal | Age | Sex | Surgery | LH or FSH;eYFP |
| 1 | <=7 w-old | XX | Ax | 0 |
| 2 | <=7 w-old | XX | Ax | 1 |
| 1 | <=7 w-old | XY | Ax | 0 |
| 2 | <=7 w-old | XY | Ax | 0 |
| Animal | Age | Sex | Surgery | GH;eYFP |
| 1 | <=7 w-old | XX | Gx | 1 |
| 2 | <=7 w-old | XX | Gx | 1 |
| 1 | <=7 w-old | XY | Gx | 2 |
| 2 | <=7 w-old | XY | Gx | 2 |
| Animal | Age | Sex | Surgery | Prl;eYFP |
| 1 | <=7 w-old | XX | Gx | 31 |
| 2 | <=7 w-old | XX | Gx | 44 |
| 1 | <=7 w-old | XY | Gx | 1 |
| 2 | <=7 w-old | XY | Gx | 1 |
| Animal | Age | Sex | Surgery | Prl;eYFP |
| 1 | <=7 w-old | XX | None | 66 |
| 2 | <=7 w-old | XX | None | 73 |
| 3 | <=7 w-old | XX | None | 72 |

**Fig.6C countings**

Cre induction efficiency quantification. On the day of dissociation, the central part of the pituitary was kept to quantify the percentage of recombination on cleft-lining SOX9 positive cells.

Manual countings

| Animal | Sex | Age | SOX9 | SOX9;eYFP | %induction |
|---|---|---|---|---|---|
| 1 | XY | 8 w-old | 1076 | 142 | 13 |
| 2 | XY | 8 w-old | 651 | 100 | 15.3 |
| 3 | XY | 8 w-old | 972 | 141 | 14.5 |
| 4 | XY | 8 w-old | 1285 | 223 | 17.3 |
| 1 | XX | 8 w-old | 527 | 51 | 9.6 |
| 2 | XX | 8 w-old | 990 | 140 | 14 |
| 3 | XX | 8 w-old | 789 | 133 | 16.9 |
| 4 | XX | 8 w-old | 1331 | 204 | 15.3 |
| 5 | XX | 8 w-old | 1260 | 209 | 16.6 |

#### Figure 6E countings

Automated Prl;eYFP countings in adult dissociated AL Sox9CreERT2;ReYFP 10 days after induction

| Animal | Sex | Age | DAPI | Prl | eYFP | Prl;eYFP |
|---|---|---|---|---|---|---|
| 1 | XY | 8 w-old | 53433 | 8774 | 906 | 0 |
| 2 | XY | 8 w-old | 118487 | 21203 | 1507 | 1 |
| 3 | XY | 8 w-old | 93587 | 16065 | 932 | 1 |
| 4 | XY | 8 w-old | 124853 | 21223 | 1373 | 0 |
| 1 | XX | 8 w-old | 199922 | 51856 | 993 | 46 |
| 2 | XX | 8 w-old | 69807 | 20337 | 605 | 9 |
| 3 | XX | 8 w-old | 91630 | 38606 | 2101 | 23 |
| 4 | XX | 8 w-old | 197278 | 66190 | 3898 | 58 |
| 5 | XX | 8 w-old | 126823 | 46440 | 3086 | 44 |

From Fig.6E countings

| %induction | Number of Prl;eYFP if 100% induction | Estimate of Prl cells from SC (Number of Prl;eYFP/Prl if 100% induction) |
|---|---|---|
| 13 | 0 | 0 |
| 15.3 | 6 | 0.02829788 |
| 14.5 | 7 | 0.04357298 |
| 17.3 | 0 | 0 |
| 9.6 | 479 | 0.92371182 |
| 14 | 64 | 0.31469735 |
| 16.9 | 136 | 0.35227685 |
| 15.3 | 379 | 0.57259405 |
| 16.6 | 265 | 0.57062877 |

Automated Prl;eYFP countings in adult dissociated AL Sox9CreERT2;ReYFP after induction at P0

| Animal | Sex | Age | DAPI | Prl | eYFP | Prl;eYFP | %Prl;eYFP/eYFP |
|---|---|---|---|---|---|---|---|
| 1 | XY | 8 w-old | 25808 | 2986 | 189 | 11 | 5.82010582 |
| 2 | XY | 8 w-old | 20815 | 2863 | 131 | 21 | 16.0305344 |
| 3 | XY | 8 w-old | 29361 | 3890 | 195 | 8 | 4.1025641 |
| 4 | XY | 8 w-old | 39251 | 5309 | 386 | 51 | 13.2124352 |
| 1 | XX | 8 w-old | 10761 | 1002 | 66 | 6 | 9.09090909 |
| 2 | XX | 8 w-old | 20353 | 2519 | 190 | 5 | 2.63157895 |
| 3 | XX | 8 w-old | 15532 | 1914 | 214 | 10 | 4.6728972 |
| 4 | XX | 8 w-old | 24943 | 3050 | 169 | 4 | 2.36686391 |

**Fig.6E, F countings**

Counting numbers for Fig. 2, 3, 5 and 6.

**Table S5.**

| CELL.SIGNATURE | GENE |
|---|---|
| Lactotroph | Prl |
| Lactotroph | Myoc |
| Lactotroph | Drd2 |
| Lactotroph | Pou1f1 |
| Somatotrophs | Gh |
| Somatotrophs | Ghrhr |
| Somatotrophs | Pou1f1 |
| Thryotrophs | Tshb |
| Thryotrophs | Trhr |
| Thryotrophs | Dio2 |
| Thryotrophs | Cga |
| Melanotrophs | Pomc |
| Melanotrophs | Pax7 |
| Melanotrophs | Tbx19 |
| Melanotrophs | Pcsk2 |
| Corticotrophs | Crhr1 |
| Corticotrophs | Tbx19 |
| Corticotrophs | Pomc |
| Corticotrophs | Avpr1b |
| Gonadotrophs | Lhb |
| Gonadotrophs | Fshb |
| Gonadotrophs | Gnrhr |
| Gonadotrophs | Cga |
| Gonadotrophs | Nr5a1 |
| Gonadotrophs | Foxp2 |
| Endothelium | Cdh5 |
| Endothelium | Angpt2 |
| Endothelium | Pecam |
| Pericytes | Pdgfrb |
| Pericytes | Cspg4 |
| Pericytes | Acta2 |
| Macrophages | Cd68 |
| Macrophages | Cd14 |
| Macrophages | Ccr5 |
| Stem cells | Sox2 |
| Stem cells | Sox9 |
| Stem cells | Hes1 |
| Stem cells | Hey1 |
| Stem cells | Fgfr1 |
| Stem cells | Notch1 |

Cell type signatures.
